# A neuromechanical model of multiple network rhythmic pattern generators for forward locomotion in *C. elegans*

**DOI:** 10.1101/710566

**Authors:** Erick Olivares, Eduardo J. Izquierdo, Randall D. Beer

## Abstract

Multiple mechanisms contribute to the generation, propagation, and coordination of the rhythmic patterns necessary for locomotion in *Caenorhabditis elegans*. Current experiments have focused on two possibilities: pacemaker neurons and stretch-receptor feedback. Here, we focus on whether it is possible that a chain of multiple network rhythmic pattern generators in the ventral nerve cord also contribute to locomotion. We use a simulation model to search for parameters of the anatomically constrained ventral nerve cord circuit that, when embodied and situated, can drive forward locomotion on agar, in the absence of pacemaker neurons or stretch-receptor feedback. Systematic exploration of the space of possible solutions reveals that there are multiple configurations that result in locomotion that is consistent with certain aspects of the kinematics of worm locomotion on agar. Analysis of the best solutions reveals that gap junctions between different classes of motorneurons in the ventral nerve cord can play key roles in coordinating the multiple rhythmic pattern generators.

## Introduction

Understanding how behavior is generated through the interaction between an organism’s brain, its body, and its environment is one of the biggest challenges in neuroscience [10, 11, 45]. Understanding locomotion is particularly critical because it is one of the main ways that organisms use to interact with their environments. Moreover, locomotion represents a quintessential example of how behavior requires the coordination of neural, mechanical, and environmental forces. *Caenorhabditis elegans* is a particularly ideal candidate organism to study the neuromechanical basis of locomotion because of the small number of neurons in its nervous system and the reconstruction of its neural and muscle anatomy at the cellular level, which has led to a detailed map of the connectivity of its the nervous system [17, 78, 84]. However, despite the available anatomical knowledge, how the rhythmic patterns are generated and propagated along the body to produce locomotion is not yet fully understood [16, 29, 90].

As with many other organisms, there are likely multiple mechanisms, intrinsic and extrinsic to the nervous system, contributing to the generation, propagation, and coordination of rhythmic patterns that are necessary for locomotion in *C. elegans* [74]. *Until recently, the majority of experimental work on C. elegans* locomotion had been focused on understanding the role of extrinsic contributions, specifically the role of stretch-receptor feedback. The proposal that stretch-receptor feedback plays an important role in the generation of movement in the nematode dates back to the reconstruction of the connectome [84]. There has since been evidence of mechanically gated channels that modulate *C. elegans* locomotion [76, 89], as well as evidence of a direct relationship between body curvature and neural activity [83]. However, coordinated rhythmic patterns can also be produced intrinsically, while remaining open to modulation through extrinsic feedback. Intrinsic rhythmic pattern generators are known to be involved in a wide variety of behaviors in a number of different organisms, including insect flight, swimming in molluscs, gut movements in crustaceans, and swimming and respiration in vertebrates [2, 19, 32, 43, 56, 61]. In an intrinsic rhythmic pattern generator, the rhythmic pattern can be generated through the oscillatory properties of pacemaker neurons or it can emerge from the interaction of networks of non-oscillatory neurons [32]. Recent experiments have provided support for the role of intrinsic rhythmic pattern generation in *C. elegans* locomotion [27, 28, 87].

It is increasingly acknowledged that simulation models play an important role in elucidating how brain-body-environment systems produce behavior [10, 14, 36, 39, 68]. In *C. elegans*, there has been a surge of theoretical work focused on understanding the neuromechanical basis of locomotion. By taking into consideration the mechanics of the body and its interaction with the environment, several computational models have demonstrated that extrinsic pattern generation alone can result in locomotion [5, 26, 30, 40, 55, 62, 83]. There have been a handful of models that have considered the potential role of intrinsic rhythmic pattern generators in *C. elegans* locomotion [21, 22, 42, 47]. Some of these models have considered a circuit capable of intrinsic rhythmic pattern generation in the head motorneurons [40, 42] or in the command interneurons [21]. Some of the models have imposed a neural activation function in the form of a travelling sine wave to drive a mechanical body to produce movement [22, 65]. Only a few models have considered the generation of rhythmic patterns from networks of motorneurons in the ventral nerve cord [47, 64]. In this model we consider both the generation of rhythmic patterns from multiple networks of motorneurons in the ventral nerve cord and the dynamic interaction of these neural patterns with the mechanical body and environment to produce movement.

Given the focus on extrinsic contributions to the generation, propagation, and coordination of rhythmic patterns controlling locomotion in *C. elegans*, current models have left a major question unanswered: Can multiple network rhythmic pattern generators in the ventral nerve cord coordinate their activity to produce the traveling wave necessary for forward locomotion in the absence of stretch-receptor feedback? And importantly, what are the different possibilities for how this could be accomplished in the worm? In this paper, we coupled multiple repeating neural units in the ventral nerve cord (VNC), whose connectivity was derived from the *C. elegans* connectome [34], to a model of the worm’s muscular system [40] and mechanical body [6], which in turn was situated in a simulated agar environment. In order to examine the feasibility of multiple network rhythmic pattern generators to contribute to forward locomotion within the VNC, the current model deliberately leaves out stretch-receptor feedback, it does not allow for the possibility of motorneurons to be pacemaker neurons, and it does not include neurons outside of the VNC to drive the circuit. We used a real-valued evolutionary algorithm to determine values of the unknown parameters of the neural circuit that optimized the ability of the coupled neuromechanical model worm to match as best as possible the ability to locomote forward on agar. To the degree that this is possible, we will learn something about what components of the worm can recreate movement under these limited conditions. Given the unconstrained nature of many problems in biology [33, 70], instead of looking for one unique model, we ran multiple evolutionary searches as a way to explore the space of parameter configurations that could lead to the behavior. Each successful search produced a distinct set of parameter values, which led to an ensemble of models that we filtered down to those that were most consistent with the worm’s behavior. The properties of this ensemble were then analyzed to identify different possible classes of solutions. Detailed analysis of the operation of the representative exemplars suggests hypotheses for mechanisms that can contribute to the generation and propagation of rhythmic patterns for locomotion in the worm.

## Methods

In this section, we describe each of the components of the model: the physical environment, the mechanical body, and the neuromuscular system; as well as the optimization technique.

### Model

The neuromechanical model (see Fig. 1) integrates the neural unit from Olivares et al., [64], with the muscular model in Izquierdo & Beer [40], and the physical model of the body and environment in Boyle et al., [6].

**Figure 1.**
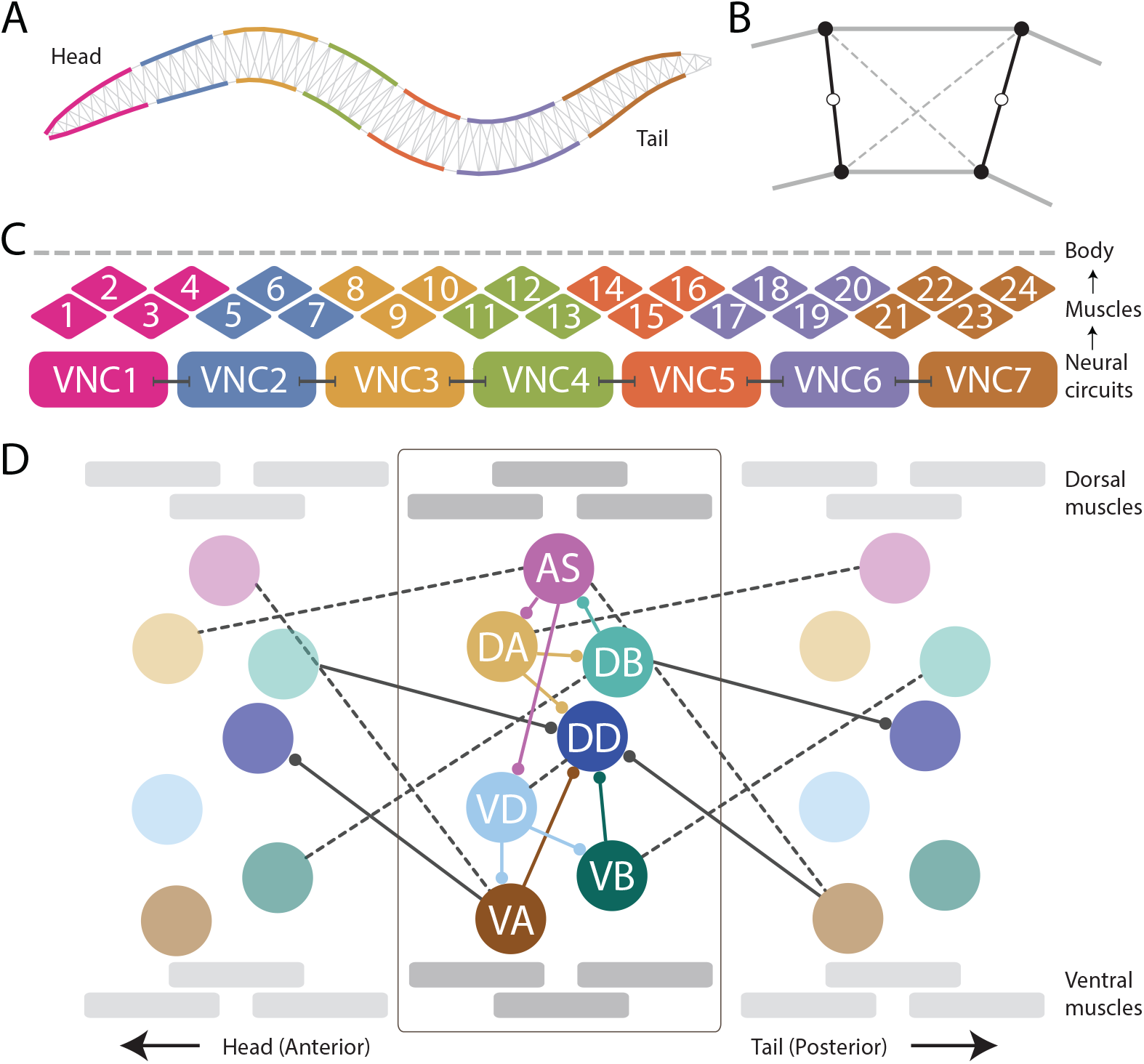
The neuromechanical model integrates a mechanical body, with a muscular system, and a repeating neural unit. [A] Following previous work [6], the mechanical body is modeled in 2D cross-section with a set of variable-width interconnected discrete segments. [B] Each segment consists of two cross-sectional rigid rods (black), damped spring lateral elements (gray), and damped spring diagonal elements (dashed). [C] Following previous work [40], muscles are modeled as elements that lie along the cuticle that can contract and relax in the dorsoventral plane, staggered along the ventral and dorsal sides of the model worm. Muscle force is distributed across all lateral elements they intersect. Seven repeating neural units innervate three ventral and and three dorsal muscles, except the most anterior and the most anterior subunits which innervate four muscles. [D] We modeled the motorneurons in the ventral nerve cord as a network composed of repeating identical subunits. The architecture of each subunit extends previous work [64] and is based on the statistical analysis of the VNC motorneurons given the missing connectome data [34]. One of seven repeating neural subunits is shown in complete detail. Intraunit connections shown in color; interunit connections shown in black. Chemical synapses shown with solid lines; gap junctions shown with dashed lines.

### Environment model

In the laboratory, *C. elegans* is typically grown and studied in petri dishes containing a layer of agar gel. The gel is firm, and worms tend to lie on the surface. The experiments in this paper focus on worm locomotion in agar. Given the low Reynolds number physics of *C. elegans* locomotion, inertial forces can be neglected and the resistive forces of the medium can be well-approximated as a linear drag *F* = −*Cv* [5, 6, 13, 62]. The tangential and normal drag coefficients for agar used in this model were taken from those reported in [4] and used in the model of the body that this work builds on [6]: *C*_‖_= 3.2 × 10^−3^ kg·*s*^−1^ and *C*_⊥_ = 128 × 10^−3^ kg·*s*^−1^, respectively [4–6, 52, 62, 80].

### Body model

When placed on an agar surface, the worm locomotes by bending only in the dorsal-ventral plane. For this reason, the worm body is modeled in 2D cross-section (Fig. 1A). The model of the mechanical body is a reimplementation of the model presented by Boyle et al., [6]. The ∼1mm long continuous body of the worm is divided into discrete segments. The width of the segments change along the length of the body as represented in Fig. 1A (for details see [6]). Each segment is bounded by two cross-sectional rigid rods (Fig. 1B). The endpoints of the rods are connected to their neighbors via damped spring lateral elements modeling the stretch resistance of the cuticle. The endpoints of the rods are also connected to the adjacent rods on the opposite side via damped spring diagonal elements modeling the compression resistance of internal pressure. The rest lengths, spring constants and damping constants of the lateral and diagonal elements are taken directly from previous work [6], which in turn estimated them from experiments with anesthetized worms [75]. The forces from the lateral and diagonal elements are summed at the endpoints of the rods and then the equations of motion are written for the center of mass of each rod. Since each rod has two translational (*x, y*) and one rotational (*ϕ*) degrees of freedom, the body model has a total of 3(*N*_*seg*_ + 1) degrees of freedom. The current model has *N*_*seg*_ = 50, so a total of 153 degrees of freedom. The full set of expressions for forces, as well as all kinematic and dynamic parameters are identical to those in previous work [5, 6].

### Muscle model

Body wall muscles in the worm are arranged as staggered pairs in four bundles around the body [1, 81]. These muscles can contract and relax in the dorsoventral plane.

Following previous work [40], muscles are modeled as elements that lie along the cuticle (Fig. 1C). The force of each muscle is distributed across all lateral elements that they intersect. Because adjacent body wall muscles overlap one another in *C. elegans*, multiple muscles can exert force on the same lateral elements. Since the model is 2D, we combine right and left bundles into a single set of 24 dorsal and 24 ventral muscles. Muscle forces are modeled as a function of muscle activation and mechanical state using simplified Hill-like force-length and force-velocity properties [35].

Following previous work [6, 40], muscle activation is modeled as a leaky integrator with a characteristic time scale (*τ*_*M*_ = 100*ms*), which agrees with response times of obliquely striated muscle [60]. The muscle activation is represented by the unitless variable 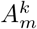 that evolves according to:

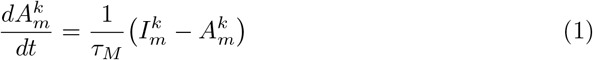

where 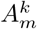 is the total current driving dorsal and ventral (*k* = {*D, V*}) muscles along the body (*m* = 1, .., 24). Also following previous modeling work [6] and experimental evidence that electrical coupling between body wall muscle cells plays only a restricted role for *C. elegans* body bend propagation [50, 83], inter-muscle electrical coupling is assumed to be too weak and therefore not included in the model.

### Ventral nerve cord circuit

As connectome data is incomplete for the ventral nerve cord [17, 78, 84], we relied on a statistical analysis of the motorneurons in relation to the position of the muscles they innervate to model the repeating neural unit along the VNC [34]. We modeled the VNC as a neural network composed of seven identical subunits (Fig. 1D). The anatomy of the repeating subunit was grounded on previous theoretical work, where we demonstrated that a subset of the components present in the statistically repeating unit found in the dataset were sufficient to generate dorsoventral rhythmic patterns [64]. The minimal configuration found in that work included motorneurons: AS, DA, DB, VD, VA, and VB; and chemical synapses: DA→ DB, DB→ AS, AS→ DA, AS→ VD, VD→ VA, and VD→VB. Given that the subunits need to coordinate their rhythmic patterns with neighboring subunits in order to produce forward locomotion, we added the following connections to adjacent neural subunits found in the statistical analysis of the VNC [34]: ASHVA^+1^, DAHAS^+1^, VBHDB^+1^, where the superscript +1 indicates that the neuron is part of the posterior subunit. We use this notation to refer to interunit connections only; for intraunit connections we leave the superscript out. The minimal configuration found in previous work [64] did not include motorneuron DD because of the lack of outgoing connections to the rest of the motorneurons within the unit, and therefore its unlikeliness to be involved in the generation of network rhythmic patterns. As the current model involves a neuromuscular system, and DD has neuromuscular junctions that allow it to drive the muscles of the worm, we included it. We also included the connections to and from DD present in the statistical analysis of the VNC [34], including intraunit connections: DA → DD, VA → DD, VB → DD, and VD → DD; and interunit connections: DB → DD^+1^, and VA^+1^ → DD.

Dorsal and ventral motorneurons in each unit drive the dorsal and ventral body wall muscles adjacent to them, respectively. The input to the body wall muscles is represented by variable 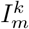 such that:

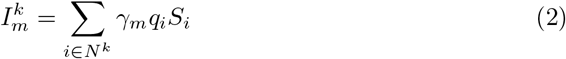

where *k* denotes whether the body wall muscle is dorsal or ventral (*k* = *D, V*) and *m* denotes the position of the muscle along the body (*m* = 1, .., 24), the set *N*^*k*^ corresponds to the dorsal/ventral motorneurons, {AS, DA, DB, DD}, {VA, VB, VD} respectively. Following previous work [6], an anterior-posterior gradient in the maximum muscle efficacy is implemented by a linearly (posteriorly) decreasing factor, *γ*_*m*_ = 0.7 (1− (((*m* − 1)*F*)*/M*)), where *γ*_*m*_ is the efficacy for neuromuscular junctions connecting motorneurons to muscle *m*, and *F* is the anterior-posterior gain in muscle contraction. *q*_*i*_ corresponds to the neuromuscular junction strength from motorneuron *i*. Finally, *S*_*i*_ corresponds to the synaptic output for each motorneuron.

The strengths of the connections in the circuit are unknown. The signs of the connections (i.e., whether they are excitatory or inhibitory) were constrained only for the neuromuscular junctions, but not for the chemical synapses between motorneurons. The AS-, A-, and B-class motorneurons are known to be cholinergic, and therefore excitatory to the muscles they innervate; D-class motorneurons are known to be GABAergic, and therefore inhibitory to the muscles they innervate [**?**, 73]. We constrained the signs of neuromuscular junctions accordingly. Note, however, we did not constrain the signs of the connections between those motorneurons and other motorneurons in the circuit because these are not known [8, 25, 51, 57, 66, 71].

### Neural model

Following studies in *C. elegans* [31, 59] and previous modeling efforts [37, 38, 40], motorneurons were modeled as nodes with simple first order nonlinear dynamics [3],

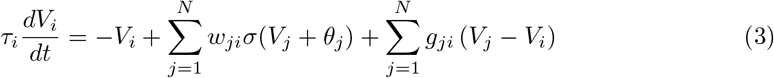

where *V*_*i*_ represents the membrane potential of the *i*^*th*^ neuron relative to its resting potential. The time-constant of the neuron is represented by *τ*_*i*_. The model assumes chemical synapses release neurotransmitter tonically and that steady-state synaptic activity is a sigmoidal function of presynaptic voltage [48, 53, 85], *σ*(*x*) = 1*/*(1 + *e*^−*x*^). *θ*_*j*_ is a bias term that shifts the range of sensitivity of the output function. The synaptic weight from neuron *j* to neuron *i* is represented by *w*_*ji*_. In line with previous theoretical work [37, 46, 85], the electrical synapses were modeled as bidirectional ohmic resistances, with *g*_*ji*_ as the conductance between cell *i* and *j* (*g*_*ji*_ *>* 0). The indices *i* and *j* used for the chemical synapses and the gap junctions represent each of the motorneurons in the circuit (AS, DA, DB, VD, VA, and VB) and the specific connectivity between them is given by the neuroanatomy (Fig. 1D). Self-connections were included in the chemical synapses term to allow for the functional equivalent of active membrane conductances which have been reported for *C. elegans* neck muscle motor neurons [31, 59]. This allows the neural model to reproduce the variety of graded activity that has been described in the free-living nematode *C. elegans* [31, 53, 54, 59]. Specifically, by changing the strength of the self-connection on each neuron, that model neuron can be either smoothly depolarized or hyperpolarized from a tonic resting potential [59], or bistable, with nonlinear transitions between a resting potential and a depolarized potential [31].

## Numerical methods

The model was implemented in C++. The neural model was solved by Forward Euler method of integration with a 0.5ms step. The body model was solved using a Semi-Implicit Backward Euler method with a 0.1ms step.

### Evolutionary algorithm

All neural circuits described in this article were produced using a simple model of evolution known as a genetic algorithm. The parameters to be searched, such as the weights and signs of the connections, were encoded as a vector of real values. A population of such vectors was maintained. Initially, the vectors in this population were randomly generated. In each generation, the fitness of every individual in the population was evaluated. A new generation of individuals was then produced by applying a set of genetic operators: selection, recombination, and mutation. Once a new population had been constructed, the fitness of each new individual was evaluated, and the entire process repeated.

A naive parameterization of our model would contain around 300 neural parameters. However, it makes little sense to work directly with such a large set of unconstrained parameters. Instead, we assumed that the parameters in each repeating VNC neural unit were identical. Altogether, the model was reduced to a total of 44 free parameters. There are 28 parameters that describe each of the 7 neuron classes and neuromuscular junctions: 7 biases, 7 time-constants, 7 self-connections, and 7 neuromuscular junctions. There are 15 parameters that describe the strength and sign (i.e., excitatory/inhibitory) of the connections: 10 weights for intraunit connections: 9 chemical synapses (AS→ DA, AS→ VD, DA→ DB, DB→ AS, VD→ VA, VD→ VB, DA→ DD, VB→ DD, VA→ DD) and one electrical synapse (VDHDD); 5 weights for interunit connections: 2 chemical synapses (VA → DD^−1^, DB → DD^+1^) and 3 gap junctions (DAHAS^+1^, VBHDB^+1^, ASHVA^+1^). One additional parameter, *F*, describes the anterior-posterior gain in muscle contraction.

To evaluate the ability of a configuration of neural parameters to produce locomotion, when embodied and situated, we created a fitness function with two components. The goal of the first component was to make a ventral nerve cord neural unit produce a rhythmic pattern. Specifically, the fitness function required that the B-class motorneurons produce a rhythmic pattern and that the frequency of the rhythmic pattern matched what has been observed for body bending in crawling worms:

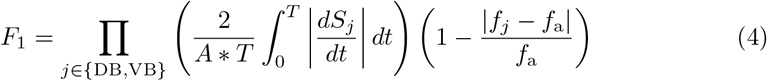

where *A* corresponds to a rhythmic pattern amplitude threshold (*A* = 0.5), *S*_*j*_ corresponds to the output of the motorneuron, *T* corresponds to the duration of the simulation, *f*_*j*_ is the frequency of neuron *j*, and *f*_a_ is the frequency of bending in the worm (*f*_a_ = 0.44Hz [12]). The first part of this equation encourages the circuit to produce a rhythmic pattern by maximizing the rate of change of neural activity in DB and VB. The contribution from this component was capped to a value of 1. The second part of the equation is aimed at matching the frequency of the worm.

The goal of the second component of the fitness function was to make the complete neuromechanical model worm move forward by matching its forward velocity to that of the worm on agar:

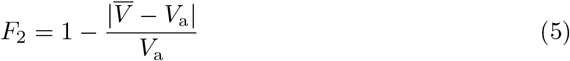

where 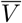 corresponds to the average velocity of the model worm over the duration of the simulation, and *V*_a_ corresponds to the average forward velocity of the worm on agar (*V*_a_ = 0.22 mm/s) [18].

## Results

### Generating an ensemble of model worms that use multiple network rhythmic pattern generators for forward locomotion on agar

As the parameters for the physiological properties of neurons and synapses involved in forward locomotion in *C. elegans* are largely unknown, we used an evolutionary algorithm to search through the space of parameters for different configurations that could produce forward movement on agar in the absence of stretch-receptors or pacemaker neurons. Because evolutionary runs with the full neuromechanical model and environment are computationally costly, we used an incremental approach. During a first stage, isolated ventral nerve cord neural units were evolved to produce a rhythmic pattern using fitness function *F*_1_ (see Methods) (Fig. 2A). Once the isolated neural units could produce rhythmic patterns (fitness *>* 0.99), they were integrated into the complete neuromechanical model and evolved to move forward in agar using a combined fitness function *F*_1_ · *F*_2_ (Fig. 2B). We ran 160 evolutionary searches with different random seeds. Of these, 104 (65%) reliably found model configurations that matched the body bending frequency and mean velocity of worms performing forward locomotion on agar (Fig. 2C). In other words, the evolutionary search consistently found configurations of the neuroanatomical circuit that could produce forward locomotion through the coordination of multiple network rhythmic pattern generators along the ventral nerve cord.

**Figure 2.**
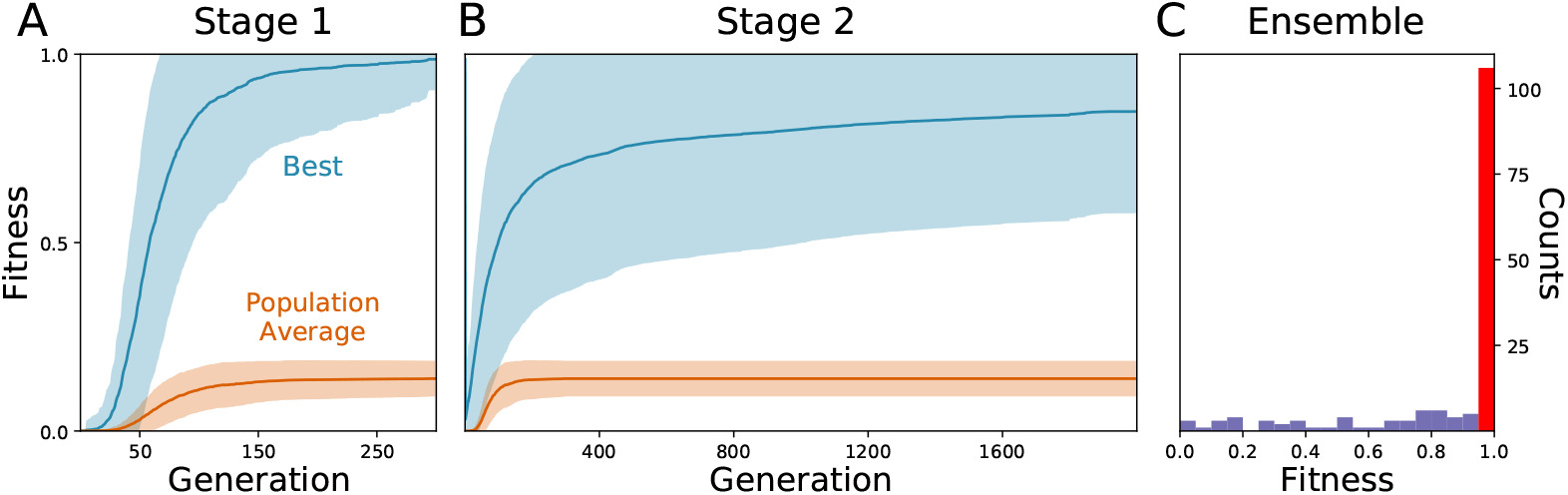
Evolutionary algorithm reliably finds model configurations that match worm forward locomotion. [A] Stage 1: Evolution of the isolated neural unit to match B-class neuron rhythmic patterns and frequency. The best and average fitness of the population are shown in blue and orange, respectively. The average over all evolutionary runs is shown in a solid trajectory and the standard deviation is shown as a lighter shade of the respective color. [B] Stage 2: Evolution of the complete neuromechanical model so as to additionally match the worm’s instantaneous velocity. Same color coding as in panel A. Distribution of best final fitness across evolutionary runs. 65% of the evolutionary searches found a solution with a fitness greater than 0.95 (red bar).

In order to focus on the subset of solutions that resembled forward locomotion in *C. elegans* most closely, we filtered the set of 104 solutions to those that matched an additional set of locomotion features that were not imposed during the evolutionary search. We applied the following three criteria: (a) Relative role of the different neuron classes in forward locomotion; (b) body curvature; and (c) trajectory curvature. Altogether, 15 solutions fulfilled all three filtering criteria (Fig. 3D). We discuss each of the criteria in turn.

**Figure 3.**
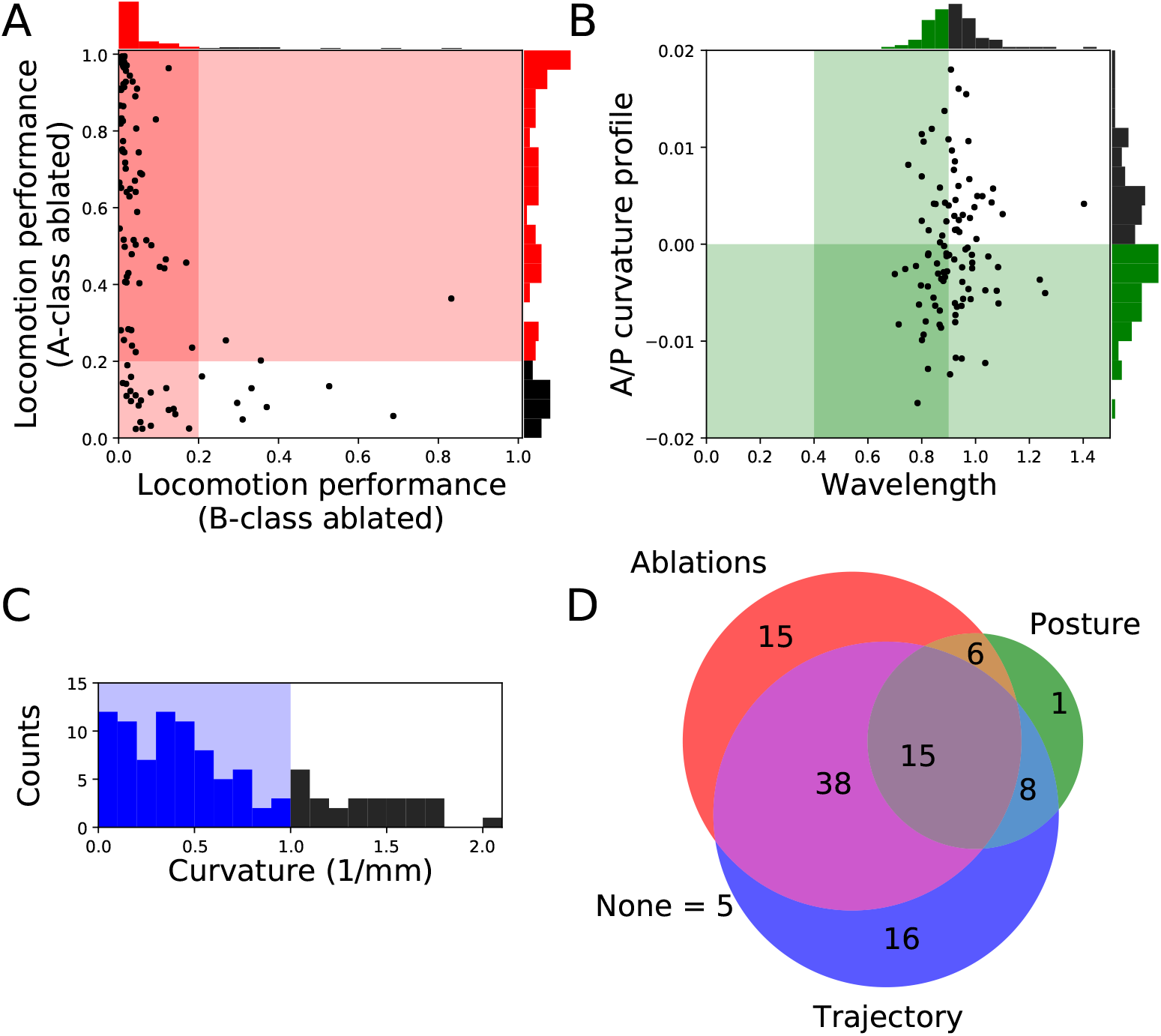
Filtering solutions to include only those that match the ranges reported in the literature about worm locomotion on agar. [A] Relative role of the different neuron classes in forward locomotion. Proportion of the forward locomotion speed maintained by each model worm when the neuromuscular junction from the B-class motorneurons (x-axis) or the A-class motorneurons (y-axis) are ablated. See *F*_2_ in Methods for the measure of locomotion performance. Each point in the figure represents a single solution. Solutions in the shaded region represent those that match the filtering criteria: (1) Ablation to A-class neuromuscular junctions should not impair forward locomotion entirely; and (2) Ablation to B-class neuromuscular junctions should impair forward locomotion performance. [B] Body curvature. Measures of the model worms’ body wavelength (x-axis) and their anterior-posterior curvature profile (y-axis) (see Supplementary Material S1 for details). Green shaded areas represent biologically plausible ranges. In the previous two figures, solutions in the darkest shaded region represent those that match both criteria. Histograms are shown for the criteria on each axis. [C] Trajectory curvature. Distribution of the radius of curvature for the trajectories of each model worm in the 2D plane (see Supplementary Material S1 for details). The blue shaded area represents solutions with a relatively straight trajectory. [D] Venn diagram representing the distribution of the 104 selected solutions according to the fulfillment of the three different filters: relative role of different neural classes in red, body curvature in green, and trajectory curvature in blue. We focus our analysis on the 15 solutions that matched all criteria.

### Relative role of the different neuron classes in forward locomotion

A- and B-class neurons have been implicated in backward and forward locomotion, respectively, through ablations performed at the larval stage, when only DA and DB neurons are present [9]. Specifically, these studies have revealed that ablating B-class motorneurons prevents forward locomotion but not backward, and that ablating A-class motorneurons prevents backward but not forward locomotion [9]. More recently, neural imaging studies in the adult have provided evidence that both A- and B-class motorneurons are active during locomotion [24]. There is also evidence that the activity of B-class motorneurons is higher during forward locomotion than the activity of A-class motorneurons, and vice-versa during backward locomotion [44]. In all evolved solutions of our model, both A- and B-class motorneurons are actively involved in forward locomotion. This is the case because the solutions are all network oscillators. So although we cannot ablate A- or B-class neurons without disrupting the network oscillation, we can silence each of their contributions to the muscles. In order to focus only on the solutions where the B-class input to muscles is necessary to produce forward locomotion but not the A-class, we simulated each solution while eliminating the neuromuscular junctions from B-class motorneurons and from A-class motorneurons, independently. We then evaluated the velocities of the model worms as a result of this manipulation (Fig. 3A). We selected solutions that met the following two criteria: (1) eliminating the A-class neuromuscular junction does not seriously compromise locomotion (i.e., velocity greater than 20% of target velocity); and (2) eliminating the B-class neuromuscular junction does compromise forward locomotion (i.e., velocity less than 20% of target velocity). A total of 74 solutions fulfilled both criteria.

### Body curvature

In addition to the frequency of the body bends, there are a number of other features of the kinematics of movement during forward locomotion that have been characterized [4, 7, 12, 18, 23, 42, 49, 69, 79, 83, 87, 88]. We further filtered our solutions based on two features: the body-bending wavelength, and the anterior-posterior curvature profile. Measurements of the wavelength of the body during locomotion in agar fall in the range of 0.4 to 0.9 body length [4, 7, 12, 18, 23, 42, 49, 69, 79, 88]. We evaluated the body wavelength in all solutions and selected those that fell within the observe range (Fig. 3B). The anterior-posterior curvature profile corresponds to the relative amount of curvature along the body axis and has been shown to be more pronounced near the head of the worm than the tail [6, 83, 87]. We evaluated the mean curvature in the anterior-posterior axis in all solutions and selected those with a negative slope in the linear regression that fit the curvature profile (Fig. 3B). Altogether, we narrowed down the 104 solutions to 30 that fulfilled both criteria (Fig. 3D).

### Trajectory curvature

The translational direction of *C. elegans* during forward locomotion tends to be relatively straight, with only a small degree of curvature in the absence of stimuli [**?**, 67]. In the evolved model worms, the straightness in the trajectory was not optimized, so the distribution of curvature in the translational trajectory is broad (Fig. 3C). In order to filter out model worms that curved much more than the worm during forward locomotion, we measured the radius of curvature for the trajectories of the centers of mass of each model worm in the 2D plane (see Supplementary Material S1 for details). We set a threshold of 1 mm in trajectory curvature radius (Fig. 3C) and we found 77 solutions that moved as straight as the worm (Fig. 3D), even in the absence of proprioceptive information.

### Behavior of model worms

Despite the absence of stretch-receptor feedback and pacemaker neurons, when simulated, all 15 selected model worms exhibited regular dorsoventral bends that propagated from head to tail (see example of one model worm in Fig. 4). The kymograph shows that the traveling wave is not perfectly smooth across the body, instead there is some amount of punctuation. This is due to a combination of simplifying factors, including that the model incorporates seven distinct neural subunits and the lack of proprioceptive feedback. It can also be seen that the traveling wave is less pronounced in the head and tail regions of the body. This is because the mechanical body itself smooths the distinct rhythmic patterns by each neural unit in relation to the unit anterior and posterior to it, which is not the case for the first and last units. The locomotion behavior of all 15 solutions can be observed in animations provided in the Supplementary Material (S2). A number of recent experiments have provided support for the possibility that multiple intrinsic rhythmic pattern generators in the ventral nerve cord could be involved in aspects of forward locomotion in the worm [27, 87]. In this section, we examine the different ways in which these model worms are consistent with some aspects of what has been observed in those experiments.

**Figure 4.**
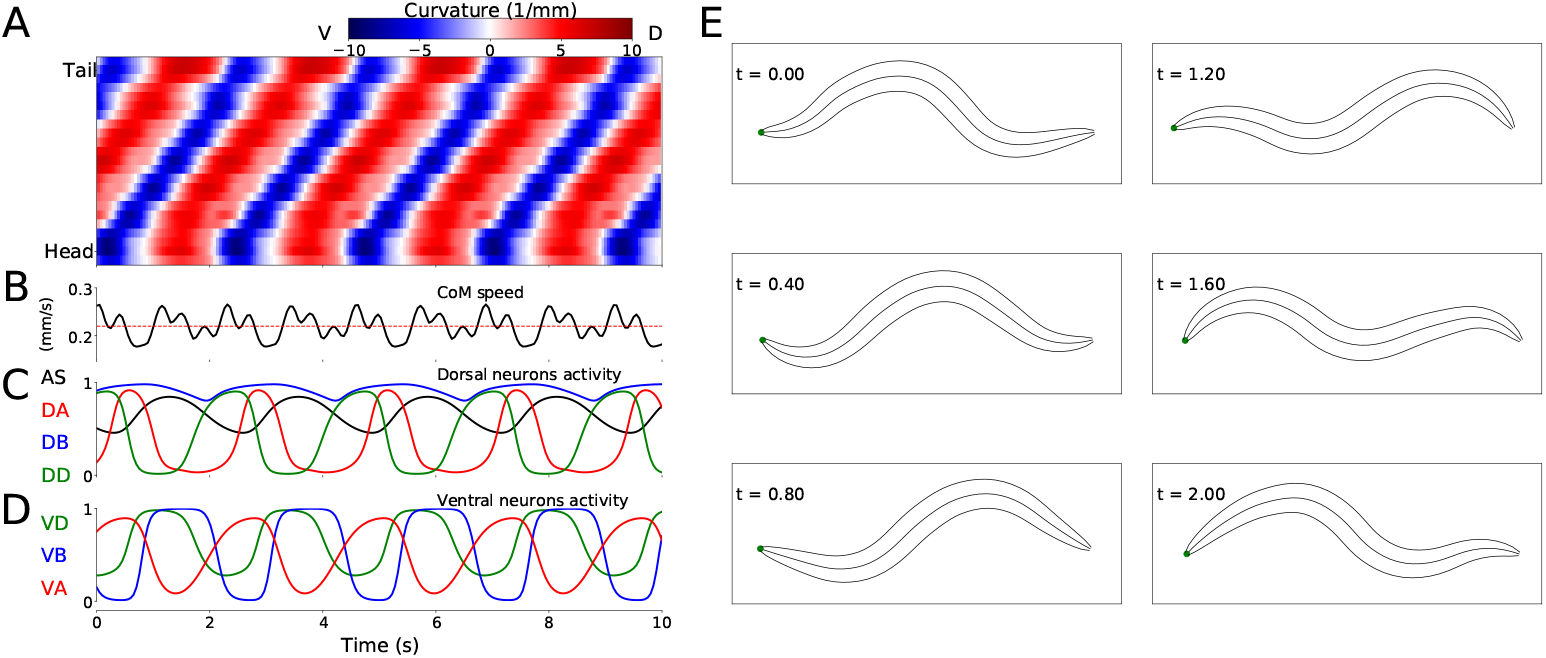
Locomotive behavior of model worms. Example from one model worm from the ensemble of solutions. [A] Kymograph depicting dorsoventral bending across the body (y-axis) and over time (x-axis). The intensity of red and blue depict dorsal and ventral curvature, respectively (see Supplementary Material S1 for the method used to calculate curvature). [B] Instantaneous velocity of the body center of mass (black trace) in relation to average velocity used for the fitness function (red line). [C] Neural activity for dorsal neurons in the first neuromuscular segment. AS in black, DA in red, DB in blue and DD in green. [D] Neural activity for ventral neurons in the first neuromuscular segment, VA in red, VB in blue and VD in green. [E] Worm body posture at different points in time (in seconds).

### Posterior rhythmic patterns persist despite anterior paralysis

Recent experiments have provided evidence that posterior dorsoventral bending persists despite anterior paralysis (Fig. 2 in [27]; and Fig. 3 and S3 in [87]). The model presented here is consistent with their experimental finding. In order to demonstrate this, we replicated the experimental condition on the model worms in two ways. First, we suppressed the neuromuscular junction activity for the three anterior-most neural units. Second, we silenced the neural activity of all neurons in those same three anterior-most neural units. Note that taking into consideration the suppression of stretch-receptor feedback was not necessary given that this model did not include stretch-receptor feedback. We examined the resulting kinematics of movement under both conditions. Specifically, we measured the magnitude of the amplitude of dorsoventral rhythmic patterns in the head and the tail. In both conditions, we observed a sharp reduction in dorsoventral bending in the head, but only a slight reduction of dorsoventral bending in the posterior regions of the body in all 15 solutions (Fig. 5A). Furthermore, coordination of the multiple rhythmic patterns in the posterior part of the body remained intact (see example from one model worm in Fig. 5B). Therefore, as with the worm, posterior dorsoventral bending persists in model worms despite anterior paralysis.

**Figure 5.**
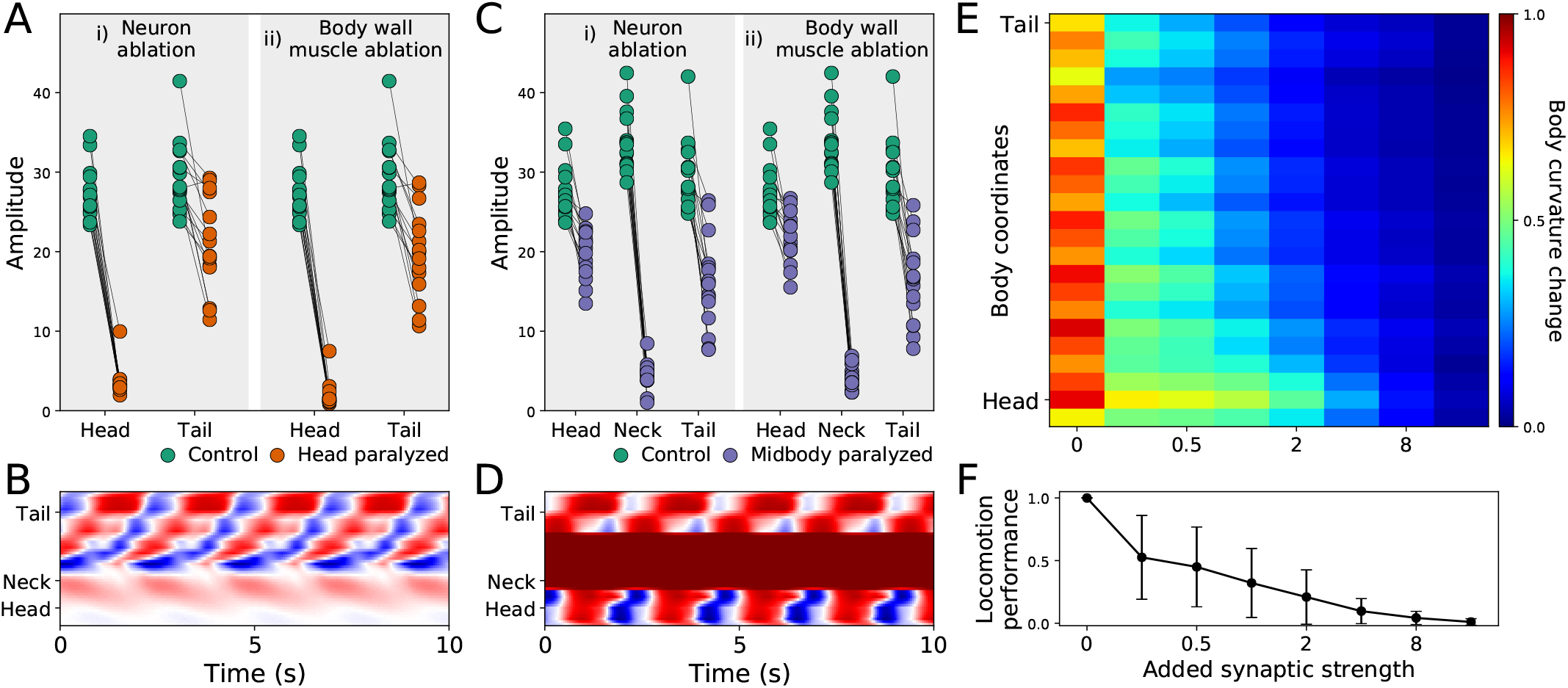
Filtered models are consistent with recent experimental observations. [A] Rhythmic posterior undulation persists despite anterior paralysis. Total bending amplitude (*y*-axis) evaluated as in previous work [27]. [i] When motorneurons in the head are ablated, bending amplitude in the head decreases but not in the tail. [ii] Similarly, when body-wall muscles (BWM) in the head are inactivated, bending amplitude in the head decreases but not in the tail. Therefore, paralysis in the head does not abolish bending in the tail. [B] Example kymograph from one model worm shows bending over time when neuromuscular junctions in the head are inactivated. The intensity of red and blue depict dorsal and ventral curvature, respectively. The color-coding is the same as the one used in Figure 4A. [C] Rhythmic undulation persists simultaneously in the head and the tail despite midbody paralysis. [i] When motorneurons in midbody are ablated, bending amplitude decreases in the midbody but not in the head or the tail. [ii] Similarly, when body-wall muscles (BWM) in the midbody are inactivated, bending amplitude decreases in the midbody but not in the head or the tail. Therefore, midbody paralysis demonstrates that head and tail are capable of simultaneous and uncoordinated rhythmic patterns. [D] Example kymograph from one model worm shows bending over time when neuromuscular junctions in the midbody are inactivated. The color-coding is the same as the one used in Figure 4A. [E] Overexpression of electrical synapses on B-class motorneurons induce complete body paralysis. Overexpression was simulated by increasing the synaptic strength of B-class gap junctions: VBHDB^+1^, DBHDB^+1^, VBHVB^+1^. [F] Speed as a function of gap junction overexpression in the model worms. The disks represent the mean and the bars represent the standard deviation. As the synaptic strength of the B-class gap junctions is increased, the speed of the model worms decreases.

### Head and tail are capable of simultaneous and uncoupled rhythmic patterns

Recent experiments have also provided evidence that the head and the tail are capable of simultaneously producing uncoupled rhythmic patterns (Fig. 3 in [27]). The model presented here is also consistent with this key component of their experimental finding. To demonstrate this, we again replicated the experimental condition in two ways. First, we suppressed the neuromuscular junction activity for the three mid-body neural units. Second, we silenced the neural activity of all neurons in those same three mid-body neural units. In both conditions, we observed that suppressing mid-body components did not eliminate body bending in either the head or the tail (Fig. 5C). In other words, like in the worm, uncoupled dorsoventral rhythmic patterns were present simultaneously in the head and the tail (see example from one model worm in Fig. 5D). It is important to note that in the experiments on the worm [27, 87], the frequency of oscillations in the anterior and posterior were different. By design, this aspect of their findings cannot be replicated by our model because every unit is identical and because neurons outside the VNC were not included. In additional experiments, we enabled and disabled each of the segments individually. We found that a single segment is not sufficient to drive bending and locomotion. Also, other than the most posterior segments, most segments contribute to forward locomotion (see Supplementary Material S6).

### Strengthening gap junctions in B-class motorneurons impairs locomotion

Finally, recent experiments have provided evidence that strengthening the gap junctions (via genetic overexpression of unc-9, one of the genes responsible for gap junctions in *C. elegans*) in the B-class motorneurons leads to constitutive paralysis in the worm [87]. Although their experiments involved the manipulation of both local electrical coupling between motorneurons as well as the descending electrical coupling from the command interneurons, the authors of that work suggest that the strong electrical couplings between the motorneurons tends to synchronize motor activity along the whole body, thus deteriorating the ability to propagate the bending wave. Our model is consistent with their interpretation of their results. To demonstrate this, we systematically increased the synaptic weight of the gap junctions interconnecting B-class motorneurons (both the VBHDB^+1^, VBHVB^+1^ and DBHDB^+1^) and measured the resulting bending along the body and speed of the model worm. As the strength of the gap junctions was increased, the bending in the body decreased. Noticeably, the effect was more pronounced in the tail than in the head (Fig. 5E). Accordingly, the velocity of the simulated worms also decreased as the strength of the gap junctions were increased, leading ultimately to total lack of movement forward (Fig. 5F). The reason for the reduced velocity is the increased synchronization of the different network oscillators along the body caused by the increased strength of the gap junctions between them. Therefore, in the model worms, strengthening the electrical couplings between the motorneurons deteriorated their ability to propagate the bending wave. There are some parallels between this result and the experiments described previously [87]. However, it is important to keep in mind the limitations of this match. First, in those experiments direct testing of the functional contribution of local electrical couplings was not possible because of experimental limitations, and therefore the electrical coupling between AVB and the B-class motorneurons could also be playing a role. Second, in those experiments stretch-receptor feedback has not been eliminated, and it could therefore also be playing a role.

### Rhythmic pattern generation and coordination in the ensemble of model worms

In the previous section we provided evidence that the simulated model worms can produce locomotion without stretch receptors and without pacemaker neurons in a way that both resembles the kinematic characterization of the worm’s forward movement on agar and is consistent with various experimental manipulations. This suggests that the way these model worms operate could be illustrative for understanding the mechanisms responsible for locomotion in the worm. Two basic mechanisms are necessary for a chain of multiple rhythmic pattern generators to drive locomotion in the worm. First, a network of neurons must be able to generate rhythmic patterns intrinsically. Second, adjacent rhythmic pattern generators must coordinate their activity with the appropriate phase delay along the anterior-posterior axis. In what follows, we examine the model worms in detail to answer the following three questions: How dominant are inhibitory or excitatory connections in the evolved rhythmic pattern generators? How do the model worms generate dorsoventral rhythmic patterns? And how do they coordinate these rhythmic patterns across the length of the body to generate a propagating wave capable of producing thrust against an agar surface?

### Excitatory-inhibitory pattern in the ensemble of model worms

It has been proposed that generating intrinsic network rhythmic patterns is difficult because the network would have to rely extensively on inhibitory connections [15]. However, the evolutionary search revealed multiple instantiations of possibilities over a wide range of the inhibition/excitation spectrum (see Supplementary Material S3 for the full set of parameters for each of the solutions). Seven out of the 15 solutions contained a majority of excitatory synapses. Furthermore, across the 15 models analyzed, it is possible to find one with any of its six chemical synapses in an excitatory configuration. This suggests a wide range of possibilities for the feasibility of multiple intrinsic network rhythmic pattern generators in the ventral nerve cord.

### Model worms use the dorsal AS-DA-DB subcircuit to generate rhythmic patterns

How do these model worms generate rhythmic patterns? To answer this question, we first determined which set of neurons are involved in producing rhythmic patterns. For a subcircuit to be capable of generating rhythmic patterns in the absence of pacemaker neurons, a recurrently connected set of neurons are required. There are three possible subcircuits in the VNC unit that are capable of generating intrinsic network rhythmic patterns: AS-DA-DB, VD-VA-DD, and VD-VB-DD (Fig. 6A). We examined whether each of these subcircuits alone could produce rhythmic patterns (Fig. 6B). We measured the total cumulative change in neural activity as an indicator of rhythmic patterns. Because the subcircuits were evaluated in isolation from the rest of the network, we examined each of them with a wide range of compensatory tonic input to each neuron. Consistent with our previous work in the isolated neural unit [64], all model worms generated rhythmic patterns in the AS-DA-DB subcircuit. Out of the 15 model worms examined, only one of them (solution M11) generated rhythmic patterns also in the VD-VB-DD subcircuit and only one (solution M6) generated weak rhythmic patterns in the VD-VA-DD subcircuit.

**Figure 6.**
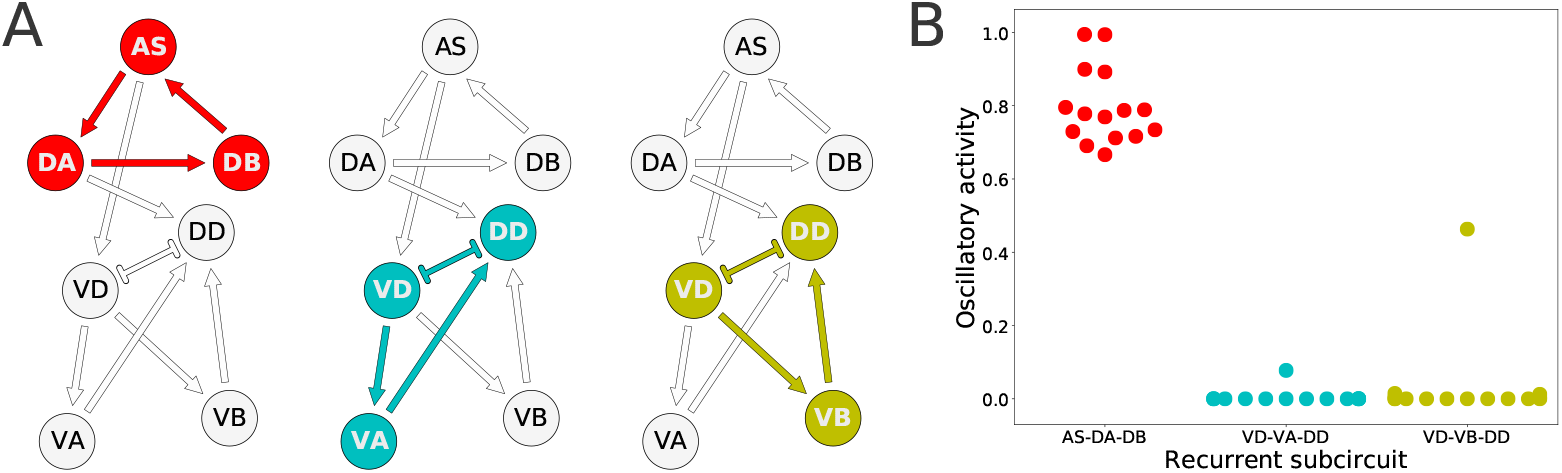
Rhythmic patterns originate primarily in the dorsal core subcircuit: AS-DA-DB. [A] Subcircuits within a neural unit where network pattern generation is possible. [B] Ability to produce rhythmic patterns for the three subcircuits in each of the solutions from the ensemble. The ability to produce a rhythmic pattern was estimated using the average of the absolute value of the derivative of the neural outputs over time, normalized to run between 0 and 1.

### Model worms use the AS →VD connection to propagate the rhythmic patterns to the ventral motorneurons

Despite the primary role of the dorsal motorneurons in the generation of rhythmic patterns, all model worms show rhythmic patterns in their ventral neural traces. In the majority of model worms (11 out of 15), the ventral motorneurons in an isolated subunit can produce rhythmic patterns (Fig. 7A). How does the rhythmic pattern propagate to the ventral motorneurons in these model worms? There are two possibilities: the AS →VD or the DA →DD chemical synapses. We examined whether the ventral B-class motorneuron could produce rhythmic patterns when either of those connections was ablated (Fig. 7). In 10 out of the 11 solutions the AS →VD (and not the DA →DD) synapse was necessary to propagate the rhythmic patterns from the dorsal core (Fig. 7B,C). This is consistent with our previous work in the isolated neural unit [64]. In one of the model worms (solution M11), neither of the connections were necessary. Recall from the previous section that this solution was the only one that also generated strong rhythmic patterns in the ventral side. There are four solutions where the ventral motorneurons do not exhibit a rhythmic pattern in the isolated subunit. In these solutions, the rhythmic pattern in the ventral motorneurons is due to interunit contributions.

**Figure 7.**
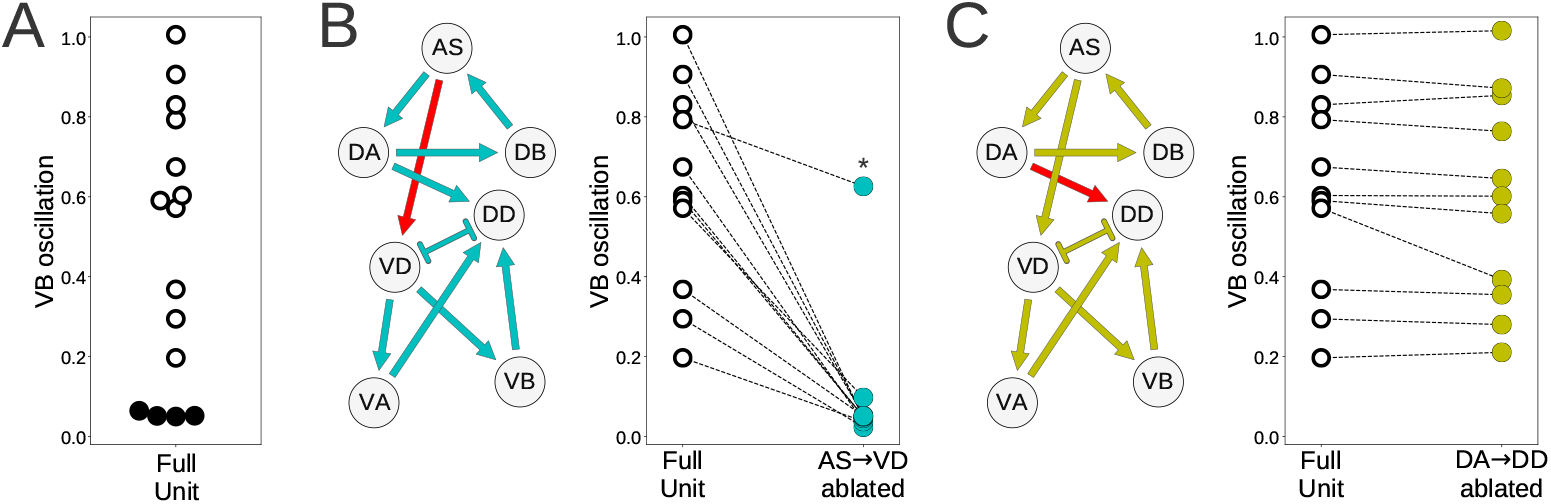
Model worms use the AS →VD connection to propagate the rhythmic pattern to the ventral motorneurons. [A] Rhythmic patterns in VB motorneurons. Four solutions do not exhibit rhythmic patterns in VB (solid black). [B] Rhythmic patterns in VB is abolished when the connection AS →VD is ablated (connection shown in red). This is true in all solutions, except M11 (shown with an asterisk), which shows rhythmic patterns in the ventral subcircuit. [C] Rhythmic patterns in VB persists when the connection DA→DD is ablated (connection shown in red).

### Rhythmic patterns coordinate through a combination of three key interunit gap junctions: AS-VA^+1^, DA-AS^+1^, and VB-DB^+1^

That the model worms move forward is evidence that the multiple rhythmic pattern generators along the body coordinate their activity. But how is the coordination between the different units achieved? To answer this question, we examined the necessity and sufficiency of each interunit connection, chemical and electrical (Fig. 8). In all the model worms examined, only the gap junctions played a role in coordinating rhythmic patterns among the different units in the VNC. The interunit chemical synapses were neither necessary nor sufficient for the coordination. In 9 of the 15 solutions examined, a single gap junction was both necessary and sufficient to coordinate the chain of multiple rhythmic pattern generators to drive locomotion forward in the worm. The VDHDB^+1^ gap junction was necessary and sufficient to coordinate rhythmic patterns in four of the solutions; the DAH AS^+1^ gap junction was necessary and sufficient in three solutions; the ASHVA^+1^ gap junction was necessary and sufficient in two. These solutions are particularly interesting because of how *simple* they are (Fig. 8). We analyze one from each of these groups in more detail in the next section. There are three solutions where multiple single gap junctions are sufficient, but no single gap junction was necessary. These solutions use *redundant* mechanisms to coordinate (Fig. 8). Finally, there are three solutions where no single connection is sufficient but several of them are necessary. These solutions are the most *complex* of the ensemble because they rely on multiple gap junctions to coordinate (Fig. 8).

**Figure 8.**
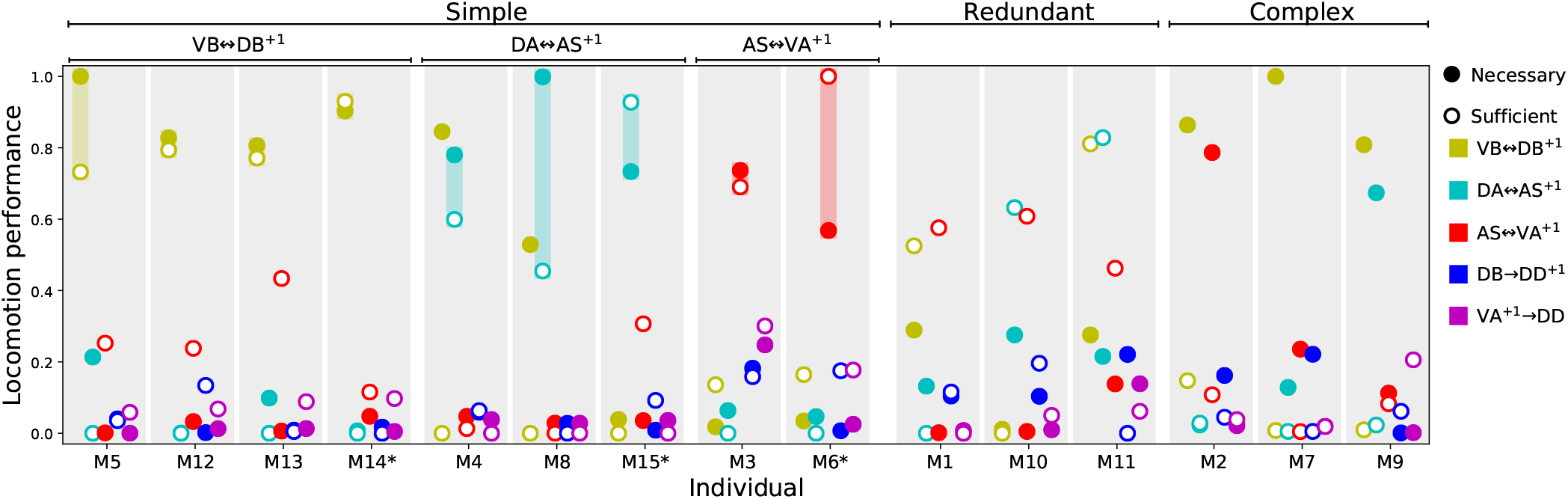
Necessity and sufficiency of each interunit connection to coordinate rhythmic patterns for locomotion in each of the model solutions. Each of the 15 solutions (labeled M1-M15) are shown on the x-axis. On the y-axis is the locomotion performance of each solution as a result of examining each of their interunit connections (labeled by color, see legend) for necessity and sufficiency. Locomotion performance was measured as the ability of model worms to match the speed of the worm (see *F*_1_ in Methods). To evaluate necessity, we ablated the connection in question and examined the worm’s ability to move forward (solid disks). To evaluate sufficiency, we ablated all but the connection in question and again examined the worm’s ability to move forward (circles). Analysis of all 15 solutions revealed three categories of strategies for coordination. “Simple” solutions correspond to those in which a single gap junction is both necessary and sufficient to coordinate the chain of multiple rhythmic pattern generators that drive locomotion. These group of solutions are further subdivided based on which of the three gap junctions is responsible for coordinating the subunits: VDHDB^+1^, DAHAS^+1^, and ASHVA^+1^. “Redundant” solutions are those in which more than one solution is sufficient to drive locomotion. “Complex” solutions are those in which no single gap junction is responsible for coordinating between units. Asterisks in the x-axis label mark the solutions with the highest single sufficient connection from each of the solutions in the “Simple” groups.

### Analysis of individual representative solutions

We have analyzed the properties of the ensemble and we have identified different possible categories of solutions based on how they coordinate the multiple rhythmic patterns. In order to understand how the circuits in the ensemble work, we need to move away from the general features of the ensemble and instead analyze in detail the operation of specific circuits. We selected three representative solutions from the “simple” group to analyze in detail, one belonging to each different cluster of solutions based on which gap junction was responsible for coordinating the subunits. Individuals were selected based on the highest performance on the sufficiency test (i.e., solution M14 for VDHDB^+1^, solution M15 for DAHAS^+1^, and solution M6 for ASHVA^+1^). Based on the results from the previous section, we simplified solutions to their minimal circuit configurations. Simulated models could still perform locomotion efficiently in these simplified configurations (Fig. 9A). In all three simplified solutions, the kinematics of movement exhibit a rhythmic pattern in the head that travels posteriorly in a way that remains consistent with what has been observed in the worm (Fig. 9B). Because all three solutions can generate movement forward, we know that the multiple rhythmic pattern generators along the body coordinate to achieve the required phase shift. From the previous section we also know that an individual synapse is sufficient to coordinate the rhythmic patterns. In this section, we examine how the coordinated phase-shift is achieved in each of these solutions.

**Figure 9.**
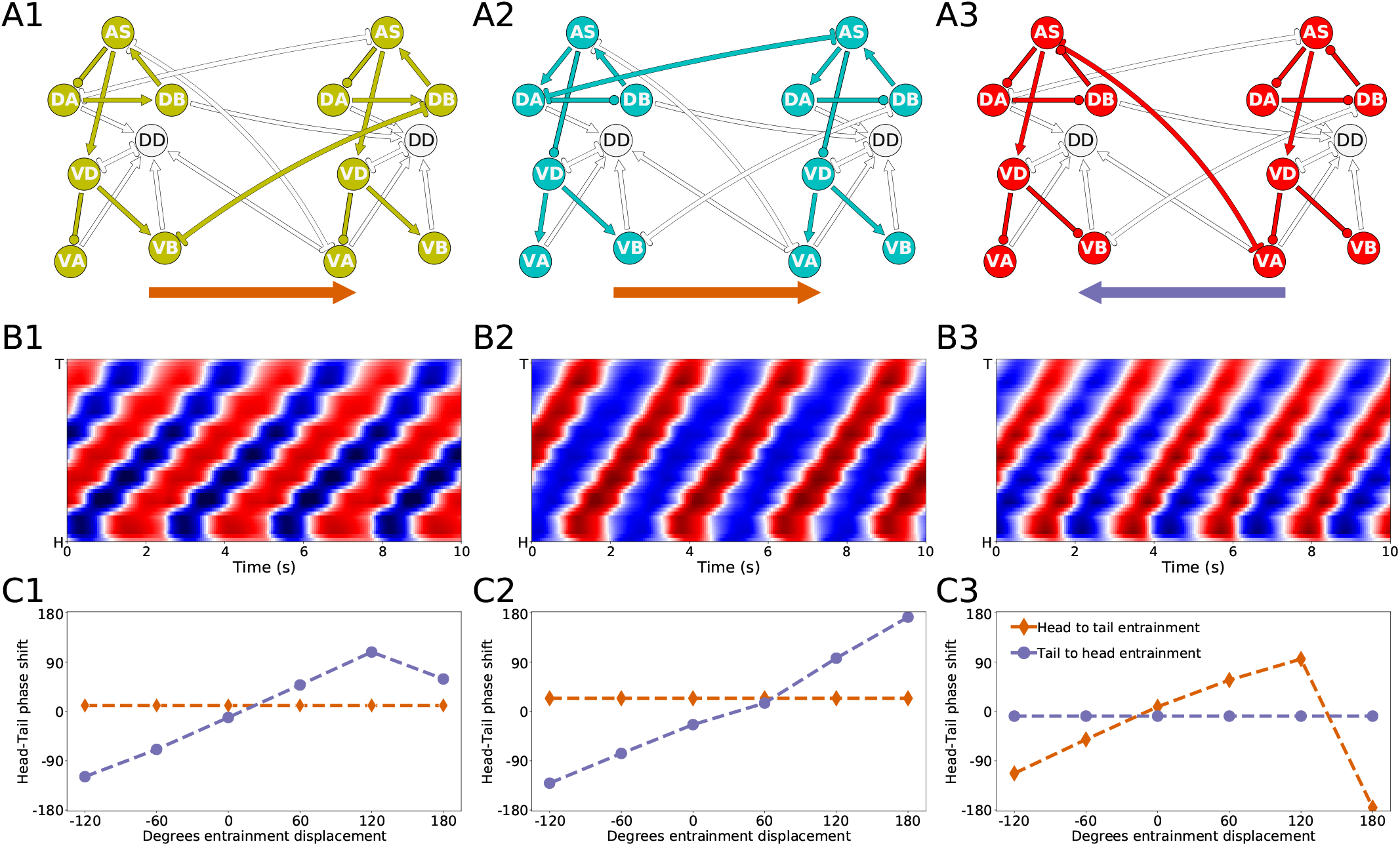
Mechanisms of anterior-posterior coordination. [A] Minimal network capable of driving locomotion in each of the solutions from the “simple” group: M14 for VDHDB^+1^ [A1], M15 for DAHAS^+1^ [A2], and M6 for ASHVA^+1^) [A3]. Arrows represent excitatory chemical synapses. Connections ending in circles represent inhibitory chemical synapses. Connections with line endings represent gap junctions. [B] Kymographs for each of the minimal configurations above show coordinated bending waves through the body. The intensity of red and blue depict dorsal and ventral curvature, respectively. The color-coding is the same as the one used in Figure 4A. [C] Entrainment analysis for each of the solutions reveals the directionality of the coordination among the subunit rhythmic pattern generators. The purple trajectory depicts the shift in phase that occurs in the posterior-most unit when the phase of the anterior-most unit is displaced. The brown trajectory depicts the shift in phase that occurs in the anterior-most unit when the phase of the posterior-most unit is displaced. In solutions M14 and M15, the anterior-most neural unit is capable of entrainning the posterior-most neural unit but not the other way around [C1, C2]. This suggests the coordination afforded by these two gap junctions is directed posteriorly. On the contrary, in solution M6, it is the posterior-most neural unit that can entrain the anterior-most unit and not the other way around [C3]. This suggests the coordination afforded by this gap junction is directed anteriorly.

### Directionality of coordination

The first thing we need to understand about coordination in these circuits is their directionality. Do anterior units influence the ones posterior to them, or vice-versa? Because the neural units along the VNC are coordinating their phases through gap junctions that allow for bi-directional communication, the directionality of coordination is not directly obvious.

First, in the solution that relies on the VBHDB^+1^ gap junction (Fig. 9A1), the anatomy suggests that the rhythmic pattern propagates posteriorly. This is because the interunit connection VDHDB^+1^ places the posterior neural subunit effectively downstream of the anterior neural subunit. The rhythmic pattern in the anterior dorsal core propagates ventrally. Then the VBHDB^+1^ gap junction coordinates the rhythmic pattern with the dorsal core unit immediately posterior to it. Therefore, in this solution, despite the bi-directionality of the coordinating gap junction, the anterior units are likely to be setting the phase of the posterior ones, and not the other way around. In order to test this hypothesis, we performed an entrainment analysis. We introduced a shift in phase first in the anterior-most neural unit and then in the posterior-most neural unit, and we measured the degree to which the rest of the neural units adopted the new phase (Fig. 9C1). As expected, when the phase was shifted in the anterior-most unit, the rest of the body adopted that shift successfully; when the phase was shifted in the posterior-most unit, the rest of the body was unaffected.

Second, in the solution that relies on the ASHVA^+1^ gap junction (Fig. 9A3), the anatomy suggests that the rhythmic pattern propagates anteriorly. This change in directionality is a result of the interunit connection ASHVA^+1^, placing the anterior rhythmic pattern generator downstream of the posterior one. The rhythmic pattern in the posterior dorsal core, once propagated ventrally, affects through the ASHVA^+1^ gap junction the rhythmic pattern of the dorsal core in the unit immediately anterior to it. Therefore, opposite to the previous model worm, in this model worm the posterior units are likely to be setting the phase of the anterior ones, and not the other way around. This is again despite the bi-directionality of the coordinating gap junction. We tested this hypothesis using the same entrainment analysis as before (Fig. 9C3). As expected, when the phase was shifted in the anterior-most unit, the rest of the body was unaffected; when the phase was shifted in the posterior-most unit, the rest of the body adopted that shift successfully.

Finally, in the solution that relies on the DAHAS^+1^ gap junction (Fig. 9A2), anatomy alone cannot tell us whether the coordination is occurring anteriorly or posteriorly. Because the connection is directly between the two neural subunits (neither one is downstream of the other), and because the coordinating component is a bi-directional gap junction, the coordination can occur in either direction. We used the same entrainment analysis as before to examine the directionality of coordination in this model worm (Fig. 9C2). When the phase was shifted in the anterior-most unit, the rest of the body adopted that shift successfully; when the phase was shifted in the posterior-most unit, the rest of the body was unaffected. Thus, in this model worm the coordination of the shift occurs from head to tail.

### Interunit phase-shift

The second aspect of the coordination that is crucial to understanding how locomotion is generated is the shift in phase between adjacent neural units. In order to examine this, we identified the approximate shift in phase that occurs at every step of the way from the DB neuron in one unit to the DB neuron in the adjacent unit (Fig. 10). We selected to measure the shift in phase between adjacent B-class neurons because of their primary role in forward locomotion. Although the neural dynamics in the model correspond to periodic patterns of activity, the specific shape of each neural activity is different. Because of the differences, we cannot relate the dynamic of two neurons as merely a shift in phase (i.e., *f* (*t*) = *g*(*t* + *T*), where *f* and *g* are the dynamics of the two neurons). Nevertheless, we can approximate the shift in phase by assuming that the neurons in the model share the same rhythmic pattern frequency. This is the case particularly in the midbody subunits. For this analysis, we used units 3 and 4 to calculate the shift in phase. In order to estimate the phase of each neuron, we calculate the middle point between the maximum and minimum rate of change for one rhythmic cycle for each neuron.

**Figure 10.**
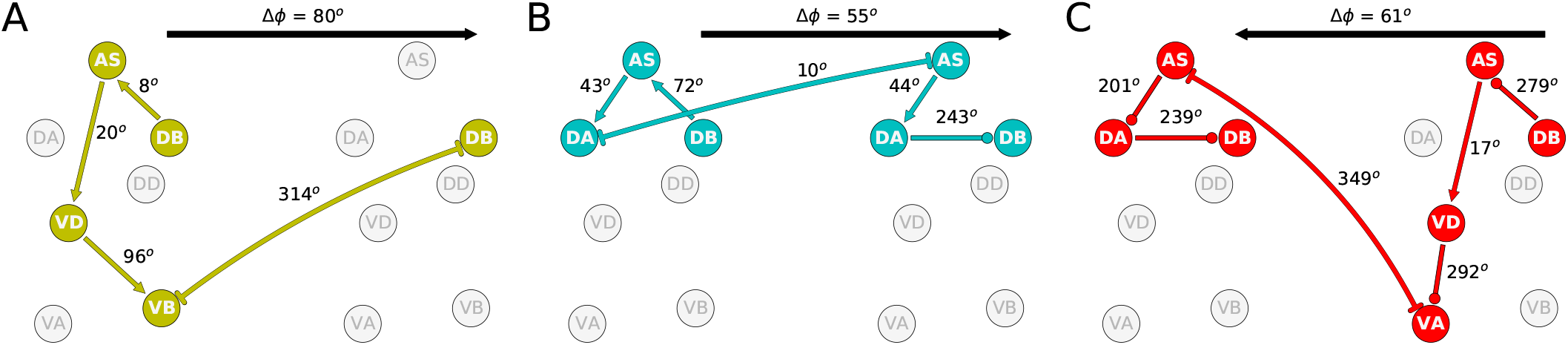
Phase delay among adjacent subunits in different networks. The highlighted neurons and connections for each network illustrates the shortest path from one DB to the DB in the adjacent neural unit in the direction of the transmission of information determined from the directionality analysis in the previous section. Despite large differences in how these different model circuits operate, the shift in phase from one unit to the one immediately posterior to it are relatively similar.

The highlighted neurons and connections for each network illustrates the shortest path from one DB to the DB in the adjacent neural unit in the direction of the transmission of information determined from the directionality analysis in the previous section. The first thing to note is that the path is different for each network (Fig. 10). Second, the shift in phase between two neurons is different among the different networks. For example, in one of the solutions (Fig. 10A), DB → AS is linked to an eight-degree shift in phase between DB and AS, whereas the same connection is linked to a 72-degree shift in phase in another network (Fig. 10B). However, despite the differences in how the model circuits operate at the level of pairwise neuron interactions, the shifts in phase from one complete neural unit to the one immediately posterior are relatively similar. It is key to note that it is this shift in phase from one unit to the next which is the primary functional activity of the network, which ultimately leads to efficient forward locomotion on agar. This analysis suggests that high level of variability at the neuron-level implementation of solutions can result in similar functional results. To highlight this result, we examined the rest of the circuits in the “simple” networks. The phase shift measured between neurons DB → AS had a mean of 154.0 degrees and a standard deviation of 109.9. Yet, adjacent units had a mean phase shift of 54.4 with a standard deviation of only 8.9.

### Interunit gap junctions present in connectome

It is important to recall that the ventral nerve cord connectome data is to date still incomplete [17, 78, 84]. Therefore, the repeating neural unit upon which we based our model is based on a statistical summary of the VNC [34]. In this section, we address how the key components that we have identified map onto the actual neuroanatomy of the worm. We examined the most recent reconstructions of the hermaphrodite and the male [17, 41, 78, 86] for the existence of the three key interunit gap junctions responsible for coordinating the multiple rhythmic patterns in the model worms. We found that all three key components occur in a large portion of the ventral nerve cord in both hermaphrodites and males (Fig 11). Moreover, because the connectome reconstruction is still incomplete in the mid-body and posterior section of the VNC, additional connections are likely to be present in these regions. The presence of additional connections can only strengthen the possibility of network oscillations. Although no single connection is present for every pair of adjacent units, there are combinations of different connections scattered throughout the full length of the ventral nerve cord. As observed in many of the evolved solutions, these three interunit gap junctions can work together to help coordinate rhythmic patterns. One thing to keep in mind is that not all of these connections are directed in the way we have idealized in the model. For example, although most ASHVA connections are directed posteriorly (i.e., with AS anterior to VA), in one of the connections the directionality is inverted (see posterior of the hermaphrodite). Assuming the directionality shown in the model, VBHDB and DAHAS will coordinate units posteriorly, and ASHVA will coordinate units anteriorly. Variability in the directionality in the connectome will have the effect of inverting the directionality of the coordination for those connections. Altogether, from these results, it seems plausible that multiple network rhythmic pattern generators in the ventral nerve cord could robustly coordinate their phases, even in the absence of stretch-receptor information, both anteriorly and posteriorly in the worm.

**Figure 11.**
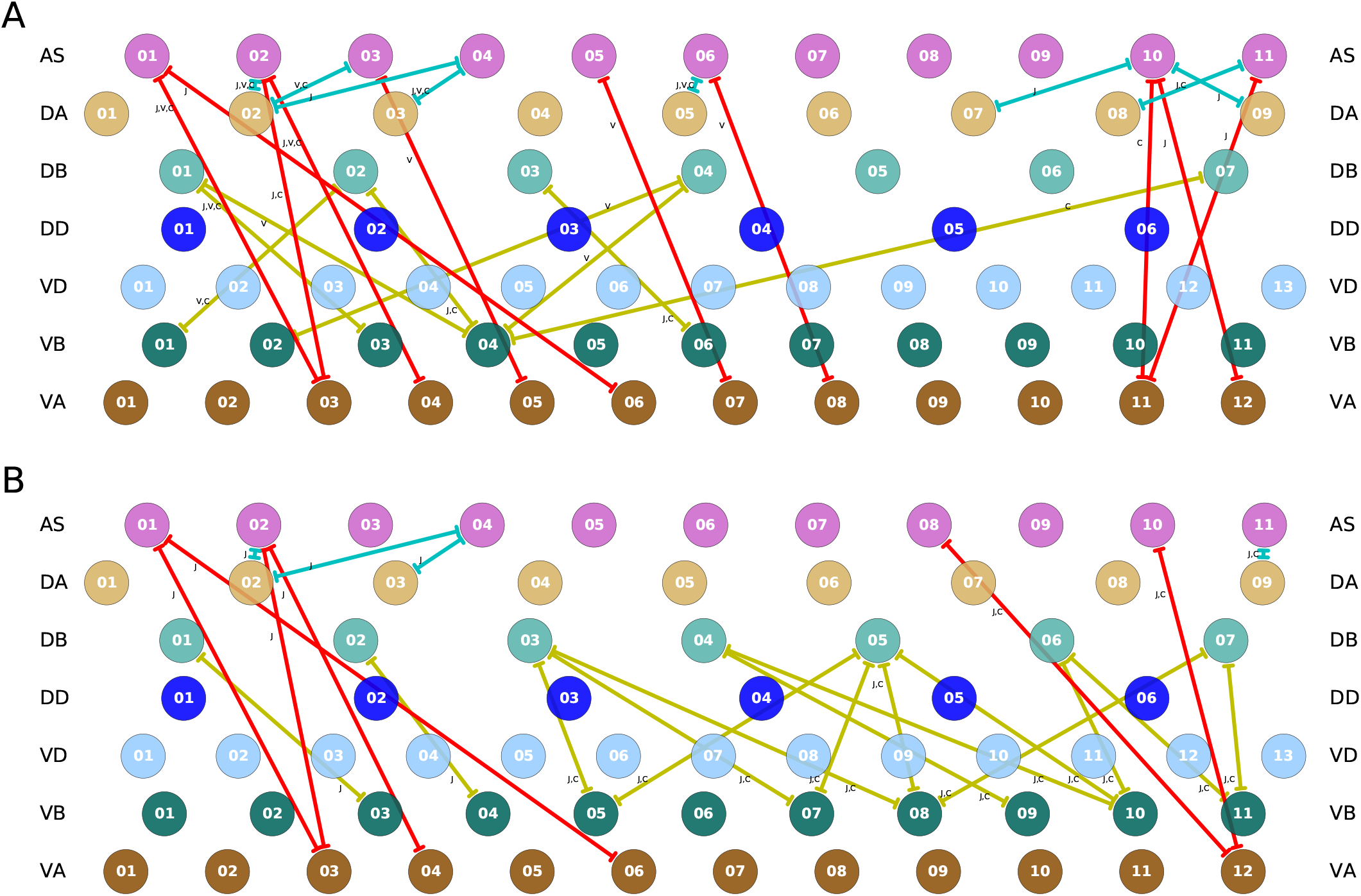
Key interunit gap junctions present in the *C. elegans* connectome for the hermaphrodite [A] and the male [B]. Gap junctions are color coded based on the pair of motorneurons they connect, as in previous figures: VBHDB in yellow, DAHAS in green, and ASHVA in red. Connections shown are present in one or several of the main datasets. For the hermaphrodite, each connection is marked as appearing in one or more of the following three datasets: j [41], v [78], or c [17]. For the male, each connection is marked as appearing in one or more of the following two datasets: j [41], or c [17].

## Discussion

Given the importance of locomotion for the worm, there is a likelihood that the worm has multiple, redundant, and overlapping mechanisms for generating and coordinating rhythmic patterns. One of the mechanisms for which there is a growing consensus is proprioception [6, 12, 26, 40, 55, 76, 83]. In this paper, we examined the possibility that multiple intrinsic network rhythmic pattern generators could coordinate through motorneuron gap junctions to produce a traveling wave along the body. The model did not aim to reproduce any one existing experiment in the literature. Instead, the model aimed at proposing a new experiment that has not been possible experimentally yet: to eliminate proprioception in the worm, to eliminate descending inputs from command interneurons, to eliminate the ability for any motorneurons to produce pacemaking activity, and to examine the worm’s ability to propagate a bending wave. The hypothesis that this model makes is that it is theoretically possible for the generation and coordination of oscillatory patterns that can generate a traveling wave, even under those conditions. If this mechanism were to be operational in the worm, then it is likely that it would be an additional contributing mechanism for forward locomotion in the worm.

We developed a model of a repeating ventral nerve cord neural circuit, and embodied and situated it to produce locomotion. In order to test whether coupling between motorneurons could be capable of generating and coordinating rhythmic patterns, the model deliberately excluded proprioceptive feedback, it excluded the ability of A- and B-type motorneurons to exhibit intrinsic oscillatory activity, and it excluded descending signals from the command interneurons. Using an evolutionary algorithm, we found 15 model configurations that reproduced the kinematics of locomotion on agar. The models were consistent with some of the recent experimental results that suggest the feasibility of multiple sources of rhythmic pattern generation [27, 87].

The multiple intrinsic oscillatory mechanism found by evolution in this model depends entirely on the collective operation of the following ventral nerve cord motor components: It depends on motorneurons AS, DA, and DB to produce the network oscillations; it depends on motorneurons VD, VA, and VB to drive ventral oscillations out of phase to the dorsal ones; and it depends on one of a set of three different inter-segmental gap junctions VB-DB, DA-AS, and AS-VA to coordinate the multiple oscillations across the length of the body. The model provides theoretical support that these components would be key places to look for intrinsic pattern generation and coordination in the worm, if it were to be identified experimentally. It follows naturally that the mechanism proposed is not robust to ablations to any of those individual components. To the degree that ablations to any of those individuals components disrupts the generation and coordination of rhythmic patterns in the worm, this mechanism for generating and coordinating network oscillations is not the sole or primary mechanism responsible for forward locomotion in the worm. In the absence of any one of the VNC motorneurons neurons, this mechanism would cease to be a possible contributor to locomotion. The hypothesis that this model makes is that in the absence of stretch receptor feedback, but otherwise with all neurons intact, it is theoretically possible for the worm to generate and coordinate multiple intrinsic oscillations. This has not been tested experimentally yet. Furthermore, the model postulates that the components mentioned above would play key roles. We discuss each of them in turn.

First, the models demonstrated that rhythmic patterns can be generated in a small subcircuit within each subunit of the model. The network of neurons identified extends the circuit proposed in previous work [64], and it includes the collective participation of all motorneurons in the VNC. However, we know from experimental studies that ablation of AS motorneurons disrupts coordination of the bending wave [77]. Together, this provides further evidence to the already strong hypothesis that stretch receptor feedback is a fundamental contributor to forward locomotion. It also suggests that, if the multiple intrinsic network oscillations mechanism is possible in the worm, it is a secondary, complementary contributor to its locomotion. Furthermore, if the nematode has multiple redundant mechanisms for producing locomotion (using stretch receptor feedback; using pacemaker neurons; and using network oscillators), then it follows that eliminating any one of the components in one of the three mechanisms will not cause a full disruption. To fully test the hypothesis of network rhythmic pattern generation will require analyzing the subcircuit of a VNC neural unit in isolation from proprioceptive feedback, without descending inputs from command interneurons, and in conditions where A- and B-type motorneurons are not generating intrinsic oscillations.

Second, although recent experimental work has focused on demonstrating the importance of proprioception for coordinating the multiple rhythmic pattern generators within the ventral nerve cord [27, 28, 82, 87], the results of our model suggest that the coordination between multiple rhythmic pattern generators can be achieved by any one of the repeating interunit electrical synapses between motorneurons. Specifically, analysis of the ensemble of models suggested three candidate electrical junctions: VBHDB, DAHAS, and ASHVA. These gap junctions are present throughout the ventral nerve cord in the available connectome data. However, only a complete anatomical reconstruction of the ventral nerve cord will reveal the full extent of their presence. To test this hypothesis would require examining whether stimulation of one set of motorneurons in the ventral nerve cord can entrain the anterior or posterior motorneurons in the absence of proprioceptive feedback and with descending input from command interneurons disabled.

Finally, analysis of the ensemble of models also demonstrated that coordination between the multiple intrinsic network rhythmic pattern generators can be achieved in either the anterior or posterior direction, and some solutions coordinate in both directions. Regardless of the direction of neural pattern coordination, the behavior of the integrated neuromechanical model was a posterior traveling wave that moved forward on agar. Altogether, analysis of the solutions revealed how different configurations of the network can generate and coordinate multiple rhythmic pattern generators that will result in almost identical forward locomotion behavior.

Given the deliberate effort to remove from the model a number of components that are known to play important roles in forward locomotion for the sake of addressing a question that had not been tackled yet, the model naturally reveals discrepancies with a number of experiments. Specifically, there is evidence that: silencing A-class motor neurons does not entirely disrupt forward locomotion [87]; ablating AS does not disrupt forward locomotion entirely but leads to a ventral bias in the bending [77]; and D-class motor neurons may not be playing a role in coordinating body waves under certain conditions [20, 58]. In contrast, the multiple network rhythmic pattern generators hypothesis in this model relies on the participation of all motor neurons, including classes A, AS, and D. Although there is further experimental work that remains to delineate with precision the role of each of those neurons in forward locomotion, there is one relatively trivial way to resolve the inconsistency between the model and the hypothesis that those neurons play no substantial roles in forward locomotion: the re-introduction of proprioceptive feedback back into the model as an independent mechanism for generating reflexive rhythmic patterns. With such a mechanism in place, silencing those neuronal classes would no longer cause disruptions to forward locomotion, as evidenced by several models that do not include those neurons, do include proprioception, and generate forward locomotion [6, 40]. That proprioceptive feedback has been removed from the model in this paper should not be taken as a postulation that it plays no role. On the contrary, we acknowledge the important role proprioceptive plays in locomotion in the worm and proceeded deliberately with the development of a model that could test the limits of whether locomotion would be possible in its absence. A second way to resolve some of the inconsistencies could arise from a complete map of the ventral nerve cord connectome. The present model only included the statistically repeating connections in the partially mapped connectome. With new connections added to the model, the possibilities for generating network rhythmic patterns only increases. With such an increase, the possibility that one of those alternatives may not require all the motor neurons in the ventral nerve cord also increases.

A more considerable discrepancy arises with the observation that body wall muscles of partially constrained worms display no activity (Fig. S1 in [83]). The results from those experiments suggest proprioceptive feedback is not just sufficient, but necessary for rhythmic activity in the ventral nerve cord. To the degree that proprioceptive feedback is demonstrated to be necessary for neural rhythmic activity, then the multiple network rhythmic pattern generators hypothesis must be discarded. However, despite those experiments, the possibility that the ventral nerve cord exhibits rhythmic neural patterns in the absence of proprioceptive feedback remains tenable. There is a possibility that the neural rhythmic activity generated is too weak to drive the muscles under the constrained condition. There is also a possibility that the constrained condition makes the activation of the muscles unfeasible. Although the lack of muscle activity in constrained worms provides strong evidence against intrinsic network pattern generation, it would be premature to discard the hypothesis altogether without first imaging neural activity while proprioceptive feedback is silenced.

Our model has a number of limitations. Locomotion in the worm is composed of many movements, including forward, backward, turns, and reversals [88]. A- and B- class VNC motorneurons receive descending input from command interneurons AVA and AVB, respectively, and activation from these neurons drives changes in the oscillatory patterns that lead the worm to transition between forward, backward, reversals, and turns [9, 44, 72, 87]. Furthermore, the locomotory pattern of the worm varies drastically with different mechanical loads from the viscosity of the environment [4, 23]. Although a more comprehensive model would address questions regarding all of those interrelated components of locomotion in the worm, the scope of this model was much narrower. The goal of this study was not to capture all aspects of the worm’s rich locomotory behavior; but to systematically search for the different parameter configurations that resulted in multiple intrinsic rhythmic pattern generators for forward locomotion on agar under the limited conditions where all VNC neurons are intact and stretch receptor feedback is not available. To study the effects of different mechanical loads on the worm’s movement, it is essential to include stretch-receptor feedback into the model. To study the changes between forward and backward locomotion and changes in speed, it is essential to include the descending input from the command interneurons.

Despite the breadth of knowledge available about the worm’s connectome, the signs of its connections are yet to be elucidated, and a systematic characterization of the physiological properties of the neurons and the role they play in different behaviors remains experimentally challenging, making modeling the neural basis of its behavior a massively under-constrained problem. Our approach takes this challenge seriously, generating multiple possible hypotheses for how different parameter configurations and patterns of activity cou ld lead to the observed behavior. Despite these obvious limitations, the strength of the approach is that it integrates connectomic, physiological, and behavioral data to infer candidate configurations of synaptic properties. As experiments that map neural manipulations to behavioral kinematics increases and the data becomes public and standardized, these can be used to further constrain the optimization search and thereby hone in on the space of candidate models.

Finally, it is important to note that the fitness function used in any evolutionary experiment defines a level of constraint on the ensemble of solutions analyzed. In this work, we chose to search for model worms that could match as best as possible aspects of the worm’s movement during forward locomotion on agar that have been characterized in some detail, such as the speed of movement [12, 18]. However, the conditions of the model worm are different from those of the wild type worm, crucially no stretch receptor feedback and no information from the command input neurons. The motivation of this simplification was that to the degree that a model worm without those crucial components could replicate some aspects of the wild-type behavior, then analysis of those solutions will reveal new insights about mechanisms for producing the behavior. Future work will involve setting up different levels of constraints on the model, which will result in different ensembles, each of which might be useful to understand. One interesting avenue of research can be aimed at understanding how the ensemble of solutions changes as constraints are changed.

## Data availability

The complete model, the evolutionary algorithm, and the tools of analysis are available online: https://github.com/edizquierdo/MultipleNetworkOscillator.

## Acknowledgments

This manuscript has been released as a pre-print at https://www.biorxiv.org/content/10.1101/710566v4 [63]. This material is based upon work supported by the National Science Foundation under Grant No. 1845322.

