## Supplementary Material for "A neuromechanical model of multiple network rhythmic pattern generators for forward locomotion in *C. elegans*"

Supplementary material to the manuscript
"A neuromechanical model of multiple network rhythmic pattern generators for forward locomotion in *C. elegans*"

Erick Olivares[1], Eduardo J. Izquierdo[1,2*], Randall D. Beer[1,2]

**1 Cognitive Science Program, Indiana University Bloomington**
**2 Luddy School of Informatics, Computing, and Engineering, Indiana University Bloomington**

*****

### Contents

1. Method to estimate wavelength, trajectory curvature, and anterior-posterior curvature profile
2. Animations of forward locomotion for the ensemble of solutions analyzed
3. Parameters for the ensemble of solutions analyzed
4. Parameter comparison to Olivares et al., 2018
5. Method to estimate inter-unit phase shift and neuron phase delay
6. Contribution from individual neuromuscular segments to locomotion performance
7. Solution robustness to added synapses and interneuron inputs

### S1: Method to estimate wavelength, trajectory curvature, and anterior-posterior curvature profile

#### Wavelength

In order to estimate the wavelength of the model worm during forward locomotion, we identified the points in which the body transitioned from ventral to dorsal bending (white areas in the kymograph, Fig. 1A). We then performed a linear regression on those points across the length of the body (dashed black lines, Fig. 1A). We used this line as an estimate of the body curvature wave in time vs. body axis plane. We estimated the period of the rhythmic pattern as the horizontal separation between two consecutive regressions (yellow horizontal line, Fig. 1B). Finally, we estimated the wavelength as the vertical distance between two consecutive linear regressions (yellow vertical line, Fig. 1B).

#### Trajectory curvature

In order to estimate the curvature of the trajectory of the model worm during forward locomotion, we ran the simulation for long enough that the worm circled back to its original starting position (Fig 1C). Using the body's center of mass, we estimated the radius of the circumference generated and used this to calculate the curvature of the trajectory.

#### Anterior-posterior curvature profile

In order to estimate the anterior-posterior curvature profile of the model worm during forward locomotion, we measured the average amount of absolute bending at each region of the body, over time (curvature profiles of all model worms shown in Fig. 1D and Fig. 1E). The amount of curvature was normalized in relation to the curvature in the head for of the curvature profile for each model worm. Model worms with a positive slope showed a tendency to bend less in the head than in the tail (Fig. 1D). The opposite was true for model worms with a negative slope (Fig. 1E).

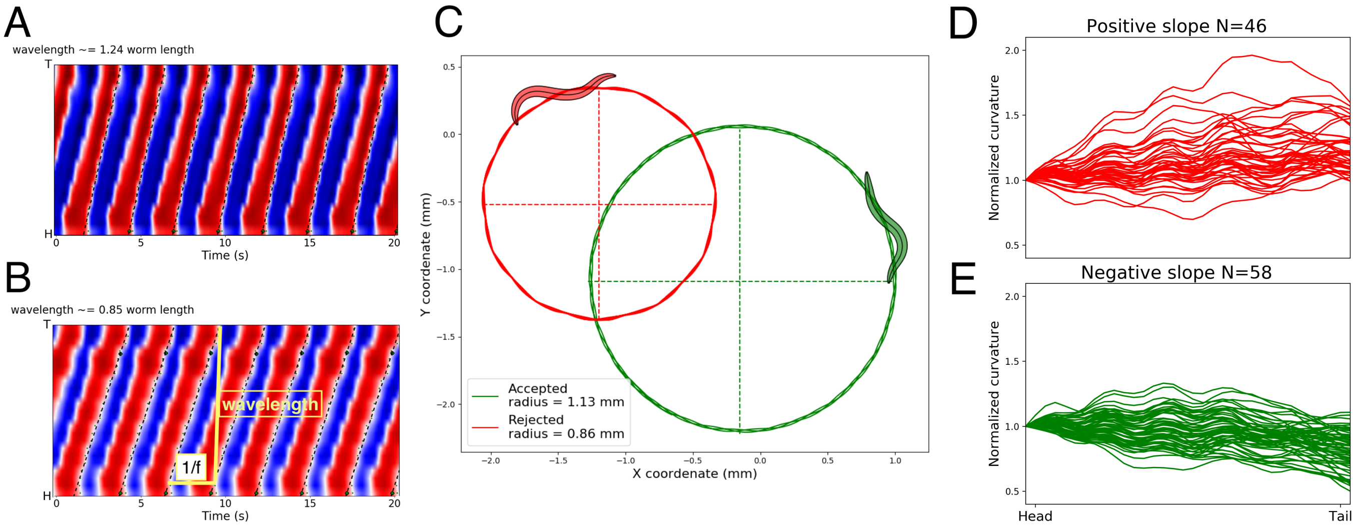


**Figure 1.** Method to estimate wavelength, trajectory curvature, and anterior-posterior curvature profile, with examples of accepted and rejected models after the filtering process. [A-B] Wavelength estimation. Example of a rejected model due to a high bending wavelength [A]. Example of an accepted model with a wavelength of 0.85, normalized to the body length [B]. [C] Trajectory curvature estimation. Example of accepted trajectory in green and rejected trajectory in red. [D-E] Anterior-posterior curvature profile estimation. The mean absolute body bending along the body axis for all model worms, with 46 solutions rejected for their trend to bend more in the tail than in the head [D] and 58 solutions with stronger bending in the head than the tail [E].

#### S2: Animations of forward locomotion for the ensemble of solutions analyzed

Each of the model worms in the ensemble was simulated on an agar environment for 25 seconds. The files for each of the model worms can be found in the Google Drive folder linked here: https://drive.google.com/open?id=14FDIk02Mbfvjapd7eSoy-yYYtuf4bckv. The videos will be added to a more permanent GitHub repository before publication. In the animations, we show the kymograph of the bending along the body over time on the top left quadrant of the screen, the neural activity in the bottom left quadrant of the screen, and we show a bird's eye view of the movement over the agar environment on the right side of the screen. We also show the shape of the worm's body in a box on the bottom right corner of the screen.

#### S3: Parameters for the ensemble of solutions analyzed

In the table below, each row represents one of the 44 parameters in the model, including the biases ($\theta_i$) for each neuron $i$, the time-constants ($\tau_i$), the self-connections ($w_i$), the chemical synapses ($w_{i \to j}$) from neuron $i$ to neuron $j$, the gap junctions ($g_{i \vdash\dashv j}$) between neuron $i$ and neuron $j$, the neuromuscular junctions ($q_i$) from neuron $i$ to its corresponding muscle, and the anterior-posterior gain ($F$). Each column represents one of the 15 solutions in the ensemble. The solutions are identified in the same way that they are referred to in the manuscript (e.g., M4, M9). Neurons with a self-connection parameter greater than 4 are bistable. Note: The neural model is not aimed at biophysical realism and the activity of the neurons does not represent voltage. Instead, the neural model is phenomenological: we are modeling the oscillations of the calcium imaging experiments. For this reason, the evolved time-constants of the neurons are relatively slow, similar to what has been observed in the calcium imaging during locomotion [3].

| | M1 | M2 | M3 | M4 | M5 | M6 | M7 | M8 |
|---|---|---|---|---|---|---|---|---|
| $\theta_{\mathrm{AS}}$ | 2.28 | -7.02 | 1.76 | 2.24 | 5.44 | 1.77 | 1.94 | -9.02 |
| $\theta_{\mathrm{DA}}$ | 14.98 | 4.80 | 7.07 | -9.51 | 12.08 | 7.20 | 0.92 | -8.78 |
| $\theta_{\mathrm{DB}}$ | 3.36 | -4.29 | 9.13 | -7.64 | 8.19 | 2.52 | -0.67 | 4.78 |
| $\theta_{\mathrm{DD}}$ | -4.73 | -9.45 | -13.66 | 3.62 | -6.38 | 5.01 | 5.45 | -6.40 |
| $\theta_{\mathrm{VD}}$ | -6.02 | 8.66 | -5.64 | 5.80 | -7.34 | -0.55 | 6.66 | 1.79 |
| $\theta_{\mathrm{VB}}$ | -0.40 | 0.85 | 9.04 | -13.13 | 11.72 | 4.35 | -2.86 | -0.57 |
| $\theta_{\mathrm{VA}}$ | -14.80 | -11.82 | 2.32 | 12.03 | 6.04 | 3.66 | -1.34 | -2.48 |
| $\tau_{\mathrm{AS}}$ $(s)$ | 0.63 | 0.26 | 0.69 | 1.30 | 1.11 | 0.25 | 0.16 | 0.18 |
| $\tau_{\mathrm{DA}}$ $(s)$ | 1.77 | 0.55 | 1.04 | 0.11 | 1.58 | 0.63 | 1.24 | 0.55 |
| $\tau_{\mathrm{DB}}$ $(s)$ | 0.10 | 0.58 | 0.10 | 0.66 | 0.15 | 0.22 | 0.15 | 0.28 |
| $\tau_{\mathrm{DD}}$ $(s)$ | 0.78 | 0.20 | 1.54 | 0.56 | 1.27 | 0.63 | 1.41 | 2.30 |
| $\tau_{\mathrm{VD}}$ $(s)$ | 0.65 | 0.91 | 1.30 | 0.32 | 0.11 | 1.26 | 0.14 | 1.05 |
| $\tau_{\mathrm{VB}}$ $(s)$ | 0.39 | 1.41 | 1.31 | 0.16 | 0.19 | 0.12 | 0.64 | 0.79 |
| $\tau_{\mathrm{VA}}$ $(s)$ | 0.98 | 0.31 | 0.18 | 0.88 | 0.38 | 0.46 | 1.15 | 2.30 |
| $w_{\mathrm{AS}}$ | 7.09 | 6.73 | 9.68 | 0.14 | 4.36 | 12.28 | -1.67 | -5.37 |
| $w_{\mathrm{DA}}$ | -14.44 | 3.74 | -0.14 | 7.82 | -10.79 | -14.69 | 9.39 | 14.37 |
| $w_{\mathrm{DB}}$ | 9.38 | 0.89 | 3.76 | 0.05 | 5.25 | 4.20 | 8.99 | 6.57 |
| $w_{\mathrm{DD}}$ | -11.75 | -0.92 | -1.82 | 1.52 | 0.78 | -6.01 | 1.60 | 6.04 |
| $w_{\mathrm{VD}}$ | -0.44 | -4.37 | -7.51 | 14.80 | -3.70 | -13.16 | -3.37 | -14.45 |
| $w_{\mathrm{VB}}$ | 4.19 | 0.99 | -9.48 | 9.15 | -10.00 | 0.34 | 4.90 | 4.98 |
| $w_{\mathrm{VA}}$ | 2.38 | 10.81 | -0.60 | 1.05 | 1.63 | 6.98 | 9.54 | -4.75 |
| $w_{\mathrm{AS}\rightarrow\mathrm{DA}}$ | -13.16 | -14.75 | -13.37 | 14.63 | -13.57 | -13.86 | -13.37 | 14.96 |
| $w_{\mathrm{AS}\rightarrow\mathrm{VD}}$ | 12.54 | -14.06 | 14.96 | 0.05 | 14.66 | 15.00 | -14.87 | 7.94 |
| $w_{\mathrm{DA}\rightarrow\mathrm{DB}}$ | -15.00 | 15.00 | -14.88 | 9.91 | -15.00 | -14.42 | -14.55 | -14.90 |
| $w_{\mathrm{DB}\rightarrow\mathrm{AS}}$ | -14.89 | 15.00 | -13.29 | -14.08 | -14.84 | -12.20 | -14.07 | 14.66 |
| $w_{\mathrm{VD}\rightarrow\mathrm{VA}}$ | -5.19 | 13.93 | -6.72 | 7.39 | -13.95 | -15.00 | 10.53 | 14.81 |
| $w_{\mathrm{VD}\rightarrow\mathrm{VB}}$ | 6.86 | -10.21 | -8.13 | 11.57 | -12.58 | -8.24 | 8.98 | 3.46 |
| $w_{\mathrm{DA}\rightarrow\mathrm{DD}}$ | 11.80 | 6.70 | -1.95 | -8.99 | 3.68 | -0.35 | 14.72 | -8.36 |
| $w_{\mathrm{VB}\rightarrow\mathrm{DD}}$ | 1.90 | -5.48 | 9.83 | -7.73 | 2.19 | -4.62 | -2.73 | 4.81 |
| $w_{\mathrm{AS}\rightarrow\mathrm{DD}}$ | -13.91 | -10.73 | 4.77 | -6.72 | 0.90 | 15.00 | 5.52 | -10.89 |
| $w_{\mathrm{DB}\rightarrow\mathrm{DD}^{+1}}$ | -7.90 | -14.18 | -14.94 | -8.14 | 9.19 | 3.43 | -5.60 | -0.26 |
| $w_{\mathrm{VA}^{+1}\rightarrow\mathrm{DD}}$ | -7.17 | -7.90 | 12.03 | 4.34 | -0.30 | 9.93 | -13.99 | 4.71 |
| $g_{\mathrm{VD}\vdash\!\dashv\mathrm{DD}}$ | 0.22 | 0.50 | 0.44 | 1.71 | 0.27 | 0.46 | 1.60 | 0.40 |
| $g_{\mathrm{AS}\vdash\!\dashv\mathrm{VA}^{+1}}$ | 0.77 | 1.76 | 0.81 | 0.96 | 0.08 | 1.41 | 0.48 | 0.02 |
| $g_{\mathrm{DA}\vdash\!\dashv\mathrm{AS}^{+1}}$ | 0.05 | 0.00 | 0.02 | 0.20 | 0.47 | 0.08 | 0.17 | 1.26 |
| $g_{\mathrm{VB}\vdash\!\dashv\mathrm{DB}^{+1}}$ | 0.53 | 1.44 | 0.84 | 0.66 | 1.18 | 0.62 | 1.03 | 0.60 |
| $q_{\mathrm{AS}}$ | 0.83 | 0.45 | 0.00 | 0.76 | 0.13 | 1.20 | 0.58 | 0.67 |
| $q_{\mathrm{DA}}$ | 0.23 | 0.63 | 0.20 | 0.51 | 0.27 | 1.13 | 0.29 | 0.00 |
| $q_{\mathrm{DB}}$ | 1.20 | 1.20 | 1.20 | 1.20 | 1.20 | 1.20 | 1.20 | 1.20 |
| $q_{\mathrm{DD}}$ | -1.10 | -0.26 | -0.90 | -0.79 | -0.10 | -0.81 | -1.20 | -0.50 |
| $q_{\mathrm{VD}}$ | -0.92 | -0.37 | -0.86 | -0.46 | -0.79 | -0.27 | -1.19 | -0.83 |
| $q_{\mathrm{VB}}$ | 1.20 | 1.20 | 1.20 | 1.20 | 1.20 | 1.20 | 1.20 | 1.20 |
| $F$ | 0.67 | 0.62 | 0.61 | 0.35 | 0.36 | 0.92 | 0.72 | 0.79 |

| | M9 | M10 | M11 | M12 | M13 | M14 | M15 |
|---|---|---|---|---|---|---|---|
| $\theta_{\mathrm{AS}}$ | -0.43 | -7.74 | -2.31 | 1.88 | 4.82 | 1.44 | -9.27 |
| $\theta_{\mathrm{DA}}$ | -4.91 | -6.31 | -8.70 | 3.37 | 4.99 | 1.86 | -9.47 |
| $\theta_{\mathrm{DB}}$ | -3.43 | 9.32 | -0.09 | -12.45 | -0.71 | -10.70 | 6.71 |
| $\theta_{\mathrm{DD}}$ | 14.99 | 3.79 | 13.36 | 0.35 | 7.84 | -11.37 | 6.33 |
| $\theta_{\mathrm{VD}}$ | -2.80 | 0.69 | -1.92 | -5.99 | 12.46 | -1.45 | 4.52 |
| $\theta_{\mathrm{VB}}$ | -0.55 | -5.89 | -7.43 | -10.16 | -4.74 | -7.51 | 4.63 |
| $\theta_{\mathrm{VA}}$ | -9.15 | -5.65 | -9.03 | -14.03 | 1.84 | -8.17 | -6.61 |
| $\tau_{\mathrm{AS}}$ $(s)$ | 0.30 | 0.30 | 1.68 | 0.29 | 1.63 | 0.49 | 0.45 |
| $\tau_{\mathrm{DA}}$ $(s)$ | 0.97 | 0.97 | 1.11 | 0.88 | 1.01 | 0.95 | 0.17 |
| $\tau_{\mathrm{DB}}$ $(s)$ | 0.21 | 0.84 | 0.12 | 0.10 | 0.11 | 0.13 | 1.11 |
| $\tau_{\mathrm{DD}}$ $(s)$ | 2.45 | 1.60 | 0.59 | 2.21 | 1.31 | 0.15 | 0.84 |
| $\tau_{\mathrm{VD}}$ $(s)$ | 0.46 | 2.46 | 0.12 | 0.10 | 0.91 | 1.28 | 0.24 |
| $\tau_{\mathrm{VB}}$ $(s)$ | 0.85 | 0.13 | 0.87 | 1.78 | 0.12 | 1.38 | 0.52 |
| $\tau_{\mathrm{VA}}$ $(s)$ | 0.54 | 0.10 | 1.02 | 0.99 | 0.35 | 1.42 | 0.10 |
| $w_{\mathrm{AS}}$ | 9.62 | 6.03 | -5.47 | -11.61 | -0.02 | -13.39 | 2.29 |
| $w_{\mathrm{DA}}$ | 0.10 | -1.35 | 0.72 | 7.15 | -0.70 | 8.14 | 10.43 |
| $w_{\mathrm{DB}}$ | -0.82 | -2.19 | 4.33 | 13.80 | 5.04 | 9.90 | 1.47 |
| $w_{\mathrm{DD}}$ | -15.00 | -8.26 | 4.87 | 12.17 | -1.53 | 7.41 | -6.94 |
| $w_{\mathrm{VD}}$ | -3.67 | -14.94 | 5.59 | 0.20 | -12.82 | -14.96 | -4.65 |
| $w_{\mathrm{VB}}$ | 5.79 | 5.48 | 5.57 | 3.77 | 2.94 | 2.40 | -8.41 |
| $w_{\mathrm{VA}}$ | -6.52 | -11.90 | -11.89 | 2.69 | -4.78 | -3.94 | 0.83 |
| $w_{\mathrm{AS}\rightarrow\mathrm{DA}}$ | 13.45 | 14.69 | 14.08 | -14.24 | -11.83 | -11.69 | 14.38 |
| $w_{\mathrm{AS}\rightarrow\mathrm{VD}}$ | 12.18 | 13.81 | 5.08 | 13.89 | -14.58 | 14.99 | -14.54 |
| $w_{\mathrm{DA}\rightarrow\mathrm{DA}}$ | 13.41 | -15.00 | -10.02 | 13.31 | -14.51 | 14.64 | -12.49 |
| $w_{\mathrm{DB}\rightarrow\mathrm{AS}}$ | -14.91 | 12.84 | 13.26 | 12.32 | -15.00 | 14.63 | 12.47 |
| $w_{\mathrm{VD}\rightarrow\mathrm{VA}}$ | -9.57 | -9.67 | -2.60 | -10.20 | 10.29 | -10.94 | 9.94 |
| $w_{\mathrm{VD}\rightarrow\mathrm{VB}}$ | -10.64 | 11.84 | 10.31 | 14.77 | 15.00 | 14.85 | 13.54 |
| $w_{\mathrm{DA}\rightarrow\mathrm{DD}}$ | 9.90 | 4.29 | -3.14 | -9.67 | -5.78 | 14.13 | -5.96 |
| $w_{\mathrm{VB}\rightarrow\mathrm{DD}}$ | -10.79 | -1.96 | -13.57 | -4.52 | -6.10 | -12.94 | 0.29 |
| $w_{\mathrm{VA}\rightarrow\mathrm{DD}}$ | -0.38 | 7.45 | -13.98 | -10.43 | -14.37 | -12.78 | -10.81 |
| $w_{\mathrm{DB}\rightarrow\mathrm{DD}^{+1}}$ | -14.94 | -14.86 | 2.10 | 1.95 | 2.06 | 9.15 | 4.54 |
| $w_{\mathrm{VA}^{+1}\rightarrow\mathrm{DD}}$ | 3.26 | -8.15 | 0.22 | -0.66 | 1.74 | -2.14 | 14.71 |
| $g_{\mathrm{VD}\dashv\vdash\mathrm{DD}}$ | 0.13 | 1.81 | 1.18 | 1.64 | 1.47 | 1.82 | 1.41 |
| $g_{\mathrm{AS}\dashv\vdash\mathrm{VA}^{+1}}$ | 1.13 | 0.30 | 0.51 | 1.86 | 0.93 | 1.03 | 0.33 |
| $g_{\mathrm{DA}\dashv\vdash\mathrm{AS}^{+1}}$ | 1.35 | 0.44 | 0.17 | 0.07 | 0.19 | 0.03 | 0.70 |
| $g_{\mathrm{VB}\dashv\vdash\mathrm{DB}^{+1}}$ | 0.99 | 0.02 | 0.34 | 1.52 | 0.62 | 1.42 | 1.19 |
| $q_{\mathrm{AS}}$ | 1.18 | 0.53 | 1.19 | 1.19 | 0.80 | 0.55 | 0.93 |
| $q_{\mathrm{DA}}$ | 0.60 | 1.20 | 0.54 | 0.29 | 0.03 | 0.00 | 0.67 |
| $q_{\mathrm{DB}}$ | 1.20 | 1.20 | 1.20 | 1.20 | 1.20 | 1.20 | 1.20 |
| $q_{\mathrm{DD}}$ | -0.35 | -0.99 | -0.96 | -0.70 | -0.93 | -0.86 | -1.16 |
| $q_{\mathrm{VD}}$ | -0.86 | -0.52 | -0.12 | -0.27 | 0.00 | -0.30 | -0.28 |
| $q_{\mathrm{VB}}$ | 1.20 | 1.20 | 1.20 | 1.20 | 1.20 | 1.20 | 1.20 |
| $q_{\mathrm{VA}}$ | 1.10 | 0.51 | 0.41 | 0.08 | 0.65 | 1.08 | 1.14 |
| $F$ | 0.56 | 0.91 | 0.39 | 0.77 | 0.79 | 0.56 | 0.76 |

#### S4: Parameter comparison to Olivares et al., 2018 [4]

In previous work [4], we evolved an individual neural segment from the ventral nerve cord to exhibit network oscillations to match patterns of neural activity related to forward and backward locomotion. In this work, we evolved seven neural segments, with additional neurons and connections, inter-connected using additional connections, and embedded it in a mechanical body and situated it in an agar environment, to produce only forward locomotion. In this section, we compare the parameters of the solutions that we examined in the previous work with the parameters of the solutions that we examined in the current work, for the subset of parameters that the two models have in common. Previous work considered 108 solutions that matched the criteria for backward and forward neural activity observed in calcium imaging experiments (green points). This work considered 15 solutions that matched a set of features in forward locomotion on agar (black points). Overall, some parameters show similar genotype distributions. However, given the differences in the conditions explained above, and the amount of variability within solutions in each of the two conditions, we do not expect a match between the two sets of solutions.

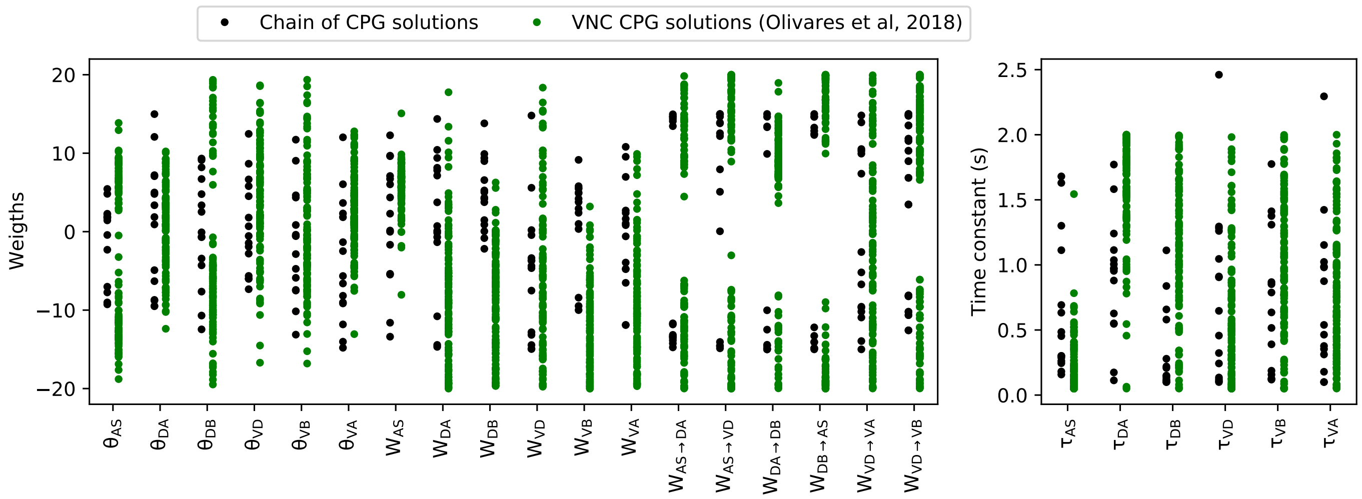


**Figure 2.** Parameters in common in selected solutions in this paper and in previous work [4].

#### S5: Method to estimate inter-unit phase shift and neuron phase delay

In order to estimate the phase shift between oscillating neurons, each neural trace was fitted by a sinusoidal function. The period was estimated by examining the derivative of the neural traces (Fig. 3), finding the maxima, and measuring the temporal distance between two consecutive maxima. The phase was estimated by assuming the midpoint between the position of the maximum and the minimum derivative of the trace as the start and end of a phase cycle (arrows in Fig 3). The phase shift between neurons was estimated by dividing the temporal difference between the two cycles by the period of the oscillation and then transforming it to degrees. We tested the performance of this ad hoc method against the performance of methods based on Fourier transforms. The method we used was more reliable than Fourier transform based methods on signals that had more than one frequency component to them.

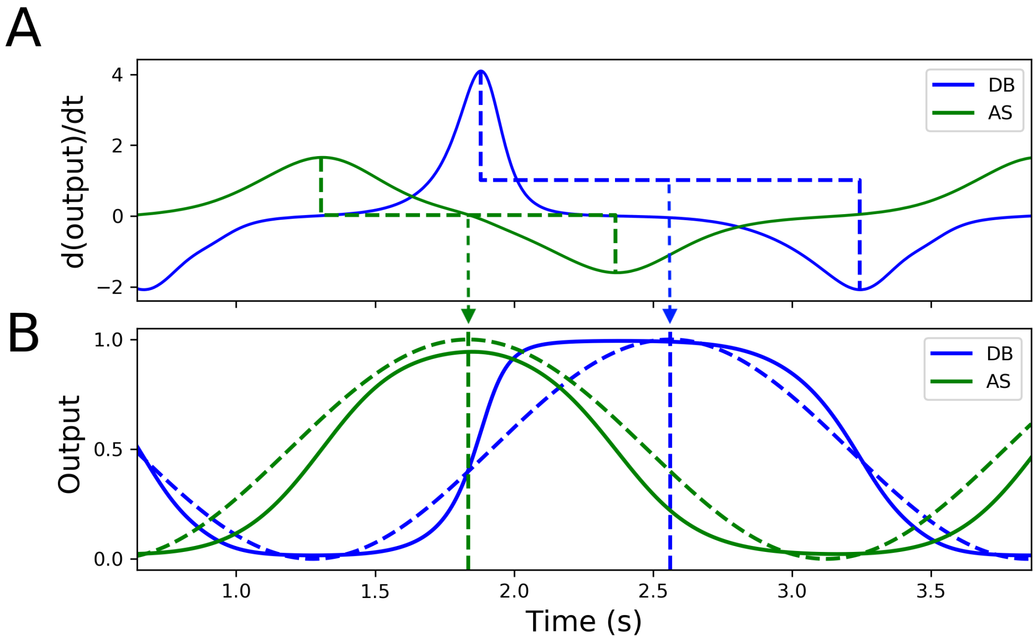


**Figure 3.** Method for estimating inter-unit phase shift. [A] The derivative of the neural traces were used to estimate the initial point of an oscillation (arrows). [B] Original neural traces (solid traces) and the sinusoidal fits (dashed traces) estimated using the derivative.

#### S6: Contribution from individual neuromuscular segments to locomotion performance

We evaluate the role of each segment in determining the speed of the model worm. Ablations of each neural segment at a time was done by fixing the neural activity of all neurons in the giving segment to a value of 0.01 (4). Anterior segments play a more relevant role in all selected models, while anterior segment relevance is more variable, disrupting locomotion in some solutions while having no effect on others. No individual segment alone is capable to move the entire worm (4), while more anterior and more posterior segments are capable to generate a minor locomotion, intermediate segments do not generate any worm displacement.

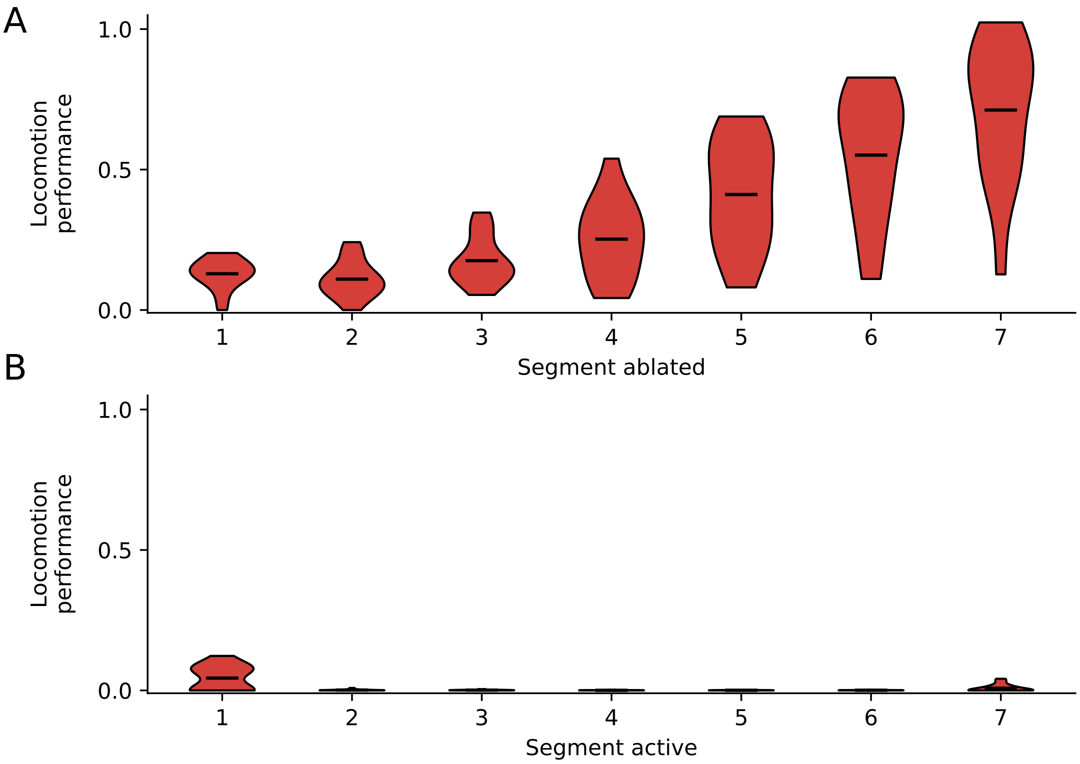


**Figure 4.** Locomotion contribution of individual segments. A) Locomotion performance distribution for the 15 selected solutions when each neural segment is ablated individually. B) Locomotion performance distribution for the 15 selected solutions when each segment is active while the rest of the segments are inactivated.

#### S7: Neuroanatomical robustness

Evolved neural circuits are unlikely to be robust to neuroanatomical changes that they have not been exposed to over their evolutionary history, particularly when no variability has been introduced during training. To introduce neuroanatomical robustness into neural circuits, the training regime can involve exposure to certain forms of neuroanatomical variability (e.g., some connections missing, some connections added, some neurons missing, some neurons added). There is a significant body of literature focused on the study of the emergence of robustness in neural circuits [2]. In this section, we included a robustness analysis of the circuit shown in Figure 4 of the main text. We added two synaptic connections (VA→VD and DD→VD) that appear in some segments of the ventral nerve cord but not all [1]. We then examined the locomotion performance as a function of a wide range of different strengths (Fig. 5). As expected, there is a gradual decay in performance as the strengths become larger. An important direction for future work in the study of the neuromechanical basis of locomotion in the worm would be to consider introducing neuroanatomical variability during evolution, and then a more thorough robustness analysis across a broad range of all possible variations.

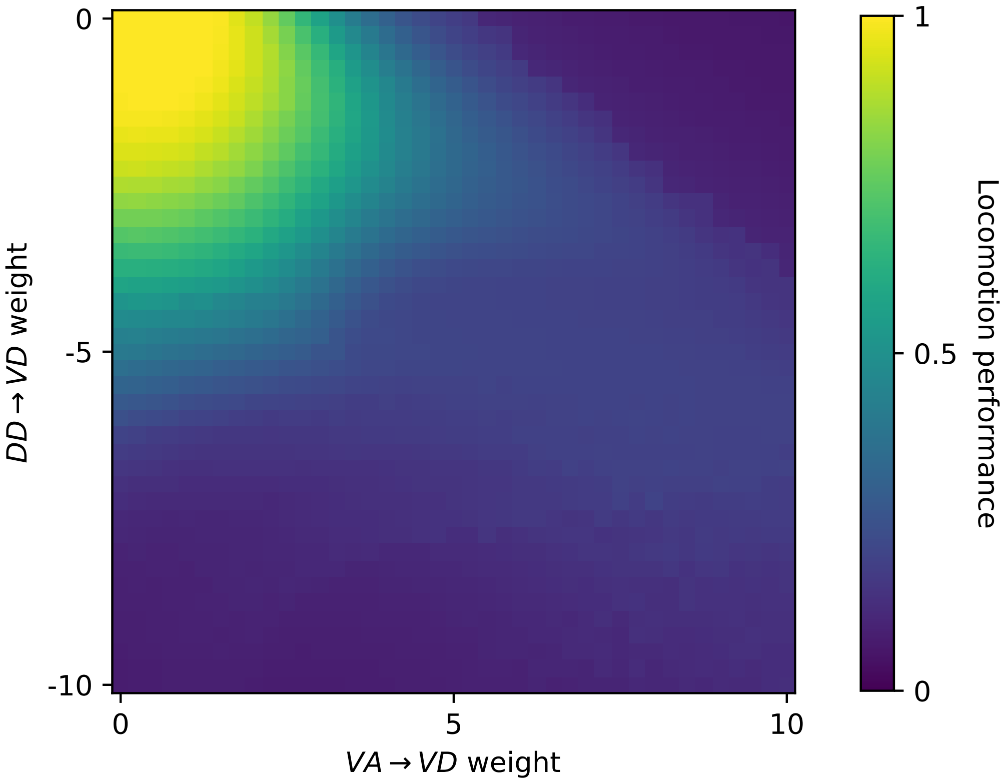


**Figure 5.** Behavioral robustness to neuroanatomical changes to the model. The analysis was performed for the circuit shown in Figure 4 of the main text. We added two synaptic connections: VA-VD and DD-VD. These connections appear in some segments of the ventral nerve cord but not all [1]. We then examined the locomotion performance as a function of a wide range of different strengths. As expected, there is a gradual decay in performance as the strengths become larger.
